# Convergence and global epidemiology of *Klebsiella pneumoniae* plasmids harbouring the *iuc*3 virulence locus

**DOI:** 10.1101/2024.01.05.574329

**Authors:** Marjorie J Gibbon, Natacha Couto, Keira Cozens, Samia Habib, Lauren Cowley, David Aanensen, Jukka Corander, Harry Thorpe, Marit AK Hetland, Davide Sassera, Cristina Merla, Marta Corbella, Carolina Ferrari, Katy ME Turner, Kwanrawee Sirikancha, Punyawee Dulyayangkul, Nour Alhusein, Nisanart Charoenlap, Visanu Thamlikitikul, Matthew B Avison, Edward J Feil

## Abstract

**Background:** *Klebsiella pneumoniae* (Kp) is an important pathogen of humans and animals, and recent reports of ‘convergent’ strains that carry both virulence and antimicrobial resistance genes (ARGs) have raised serious public health concern. The plasmid-borne *iuc* locus, encoding the siderophore aerobactin, is a key virulence factor in this species. The variant *iuc*3 is associated with porcine and human clinical isolates and is carried by mostly uncharacterised IncF plasmids.

**Methods:** We used a combination of short-read and long-read sequencing to characterise IncFIB(K)/IncFII *iuc*3-carrying plasmids harboured by 79 Kp isolates and one *K. oxytoca* isolate recovered as part of two large ‘One-Health’ studies in Italy (SpARK) and Thailand (OH-DART). Adding data from public repositories gave a combined dataset of 517 *iuc*3 isolates, and the plasmids were analysed using both clustering and phylogenetic methods.

**Findings:** We note seven large, convergent, plasmids from Thailand that have emerged through the hybridisation of co-circulating plasmids harbouring *iuc*3 and antimicrobial resistance genes (ARGs) encoding extended-spectrum beta-lactamases (ESBLs). We were also able to identify putative parental plasmids which were mostly associated with two neighbouring meat markets, as were the hybrid plasmids. Clustering and global phylogenetic analyses resolved an *iuc*3 plasmid sub-group circulating throughout Asia, with occasional examples in Europe and elsewhere. This variant carries multiple ARGs and is commonly harboured by clinical isolates, thus warranting targeted plasmid surveillance.

**Interpretation:** Our study reveals that plasmid hybridisation leading to the convergence of resistance and virulence traits may be very common, even in non-clinical (‘One-Health’) settings. Population-scale plasmid genomics makes it possible to identify putative parental plasmids, which will help to identify plasmid types that are most likely to hybridise, and what the selective consequences may be for the plasmid and host. A distinct *iuc*3 plasmid sub-variant is associated with clinical isolates in Asia which requires close monitoring.

**Research In Context:** Multiple reports of ‘convergent’ clones of *Klebsiella pneumoniae* that combine both hypervirulence and multidrug resistance (MDR-hvKp) have been published recently; a PubMed search in November 2023 using the key words ‘convergence *Klebsiella pneumoniae*’ returned 143 papers, 99 of which were published from 2020 onwards. Our study demonstrates that the hybridisation of plasmids carrying AMR and virulence genes is a frequent, ongoing, process in natural populations. The subsequent transfer of plasmids conferring both traits is thus likely to be a key driver behind the spread of convergent strains. Our study also provides an exemplar of how hybrid assemblies can facilitate large-scale global genomic plasmid epidemiology.

**Evidence before the study:** Although multiple recent reports highlight the emergence and spread of convergent Kp strains, the confluence of resistance and virulence genes within the same plasmid has not been studied at a population level, and putative parental plasmids are rarely identified. Moreover, there have been few high-resolution genomic epidemiology studies on closely related plasmids using both long and short-read data on a global scale.

**Added value:** We more than double the number of complete sequences available for plasmids harbouring *iuc*3 from 58 to 139 and provide evidence on the host lineages most likely to harbour these plasmids (e.g., ST35), and epidemiological source (e.g., pig, wild animal, human). Our comparative analysis of phylogenetic and clustering approaches will help to inform future plasmid epidemiological studies.

**Implications:** The hybridisation of plasmids harbouring virulence and resistance genes occurs frequently in natural populations, even within ‘One-Health’ settings. However, the selective drivers (if any) and evolutionary consequences of this phenomenon are unclear. There is clear utility in generating closed plasmid genomes on a population scale, and targeted plasmid surveillance on a clinical sub-variant of *iuc*3 plasmids is warranted.

## Introduction

*Klebsiella pneumoniae* (Kp) has been designated a priority pathogen by the WHO and can cause serious disease in both humans and animals^1^. Healthcare-associated Kp clones that have acquired resistance to carbapenems (CR-Kp) and/or that have become resistant to multiple other antimicrobials (MDR-Kp), are responsible for a high global public health burden^2^. These resistant clones are largely distinct from hypervirulent lineages of Kp (hvKp) responsible for severe disease in the community, including liver abscesses and pneumonia^3–6^. However, ‘convergent’ Kp lineages have emerged that combine both resistance and hypervirulence (CR-hvKP or MDR-hvKp),^7–9^. A single strain can possess both traits if virulence and resistance plasmids co-exist in the same cell, or if hybrid plasmids emerge that carry both virulence and resistance genes^10,11^. Over two-thirds of reports of phenotypically convergent isolates have been from China or South-East Asia^5,12–14^, and in particular the Chinese CR-hvKP ST11 clones are highly transmissible^14,15^. Moreover, strains, and even hybrid plasmids^16–19^, carrying both resistance and virulence traits, have also been reported in Europe, North and South America, and elsewhere^20–23^.

The *iuc* locus is a key virulence factor consisting of five genes (*iuc*ABCD and *iut*A) and encoding the siderophore aerobactin^24^. Six distinct *iuc* lineages have been identified; 1,2,2a,3,4,5, which can be further subdivided using an Aerobactin Sequence Type (AbST) scheme^25^. Whereas Kp isolates responsible for severe community-acquired infections typically carry *iuc*1 or *iuc*2 on KpVP-1 and KpVP-2 plasmids, *iuc*3 is carried by a more diverse set of IncFIB_K_-IncFII_K_ [also named IncFIB(K)-IncFII(pKP91)] plasmids. These *iuc*3 plasmids are associated with porcine isolates, but also occasionally harboured by clinical isolates^26–28^, pointing to a potential public health risk from animal reservoirs. The global diversity and epidemiology of *iuc*3 plasmids, the associated public health risk, and their possible role in disseminating resistance remains unknown. We conducted a global population analysis of plasmids harbouring *iuc*3 by combining novel hybrid genome assemblies using strains assembled during two large ‘One-Health’ surveys^26,29^ with all available public data. We found multiple examples of plasmid hybridisation leading to convergence of AMR and virulence, and the existence of an *iuc*3 plasmid sub-variant associated with clinical isolates in Asia. Targeted surveillance of this sub-variant throughout hospitals in Asia is warranted. Our study demonstrates that the ‘One-Health’ framework is pertinent for monitoring the emergence and spread of virulence, as well as resistance plasmids, and the convergence of both traits.

## Methods

### SpARK and OH-DART sampling

Details on the SpARK sample collection from Northern Italy are described in Thorpe et al., 2022^26^, and the sampling for the OH-DART project in Thailand is described in Supplementary Methods. A key difference between the studies was that the Thai isolates were selected for resistance to 3^rd^ generation cephalosporins (3GC-R) using Chromagar 3GC-R or to carbapenems using Chromagar CPE, and selected colonies were re-streaked on Brilliance Selective Medium (ThermoFisher Scientific) with either cefotaxime (2 µg/mL) or ertapenem 0.5 (µg/mL). The Italian isolates were enriched in Luria Bertani (LB) broth with amoxicillin (10 µg/ mL) and grown on SCAI media with ampicillin (10 µg/mL)^30^.

### Sequencing

Short-read sequencing of *Klebsiella* isolates from Italy has been previously described^26^. For long-read sequencing of the SpARK isolates, DNA was extracted using the Wizard DNA kit (Promega). Libraries were prepared using the rapid barcoding kit SQK-RBK004, multiplexing up to 48 isolates per run, and sequenced on a MinION or GridION device with R9.4.1 flow cells (Oxford Nanopore Technologies [ONT], Oxford, UK), using default settings. The OH-DART isolates from Thailand were sequenced externally by MicrobesNG (https://microbesng.com/), as described in Supplementary Methods. The German isolate was sequenced as described previously^28^.

### Genomic characterization and phylogenetic analysis

Genome assemblies were assigned species and multi-locus sequence types (MLST), and screened for virulence and resistance genes, using Kleborate v2.3.2 (https://github.com/katholt/Kleborate)^10^. The aerobactin locus was extracted from the assemblies and aligned using MAFFT (in Geneious 2022.1.1), after which an approximate maximum-likelihood phylogenetic tree based on a general time reversible (GTR) model was generated using FastTree v2.1.11^31,32^. A mashtree of all the Kp assemblies from Italy and Thailand was obtained using mashtree v1.2.0^33^. See Supplementary Methods for more details.

### Closed circular plasmids

Hybrid assemblies of Illumina and ONT sequence data were generated using Unicycler v0.4.8^34^. Assemblies were annotated using Prokka v1.14.6^35^. We also retrieved plasmid sequences carrying *iuc*3 from Kp from previous publications or from GenBank (Supplementary Tables 1, 2). Further details on bioinformatics analysis are provided in Supplementary Methods.

### Identification of putative parental plasmids using BLASTn

Putative parents of hybrid plasmids assigned as convergent were identified amongst the closed OH-DART circular plasmids using BLASTn (v2.14.0) with default settings. Plasmids with the longest regions of high nucleotide identity (>98%) spanning either the *iuc* locus or resistance gene loci were identified.

### Short-Read Mapping to Plasmid Sequences

We combined additional short-reads and assemblies (short-read and closed) of Kp carrying *iuc*3 (as identified using Kleborate) from the public domain with the fully closed plasmid sequences generated in this study. Snippy v4.6.0 (https://github.com/tseemann/snippy) was used to map short-reads to OH-DART_30005-KC1_2. For those *iuc*3 plasmids where only assemblies were available (and not reads), we used the Snippy contigs mode to artificially generate reads, and then treated these the same as the short-read data. These were used to generate an approximate maximum-likelihood phylogenetic tree based on a general time reversible (GTR) model using FastTree v2.1.11^31,32^. The tree was combined with metadata and output from Kleborate v2.3.2 and visualised using Microreact v251^36^.

### Statistical analysis

The statistical analysis was carried out in RStudio v2023.03.1 using R v4.3.0.

## Results

### The prevalence and provenance of *iuc*3 plasmids

We analysed genome data for a total of 517 *iuc*3 positive plasmids, and a breakdown of this dataset is given in Supplementary Table 1. Although multiple isolates from *Klebsiella* species other than Kp were sequenced in both the SpARK and OH-DART projects, *iuc*3 was exclusively found in Kp, except for a single *K. oxytoca* isolate in the SpARK data. Lineages harbouring *iuc*3 represent the full breadth of the Kp phylogeny (Supplementary Figure 1), and the ecological distribution of the *iuc*3 isolates in the SpARK and OH-DART datasets is summarised in Supplementary Figure 2. As reported^26^, *iuc*3 is strongly associated with porcine isolates in the SpARK dataset. Although only 4% of all 1705 SpARK Kp isolates were recovered directly from pigs (n=69), these account for 42/49 (85.7%) of all *iuc*3 isolates in this study. Four of the other seven *iuc*3 isolates were recovered from the pig farm environment, two from dogs and one from the urine of a hospital inpatient.

The 591 Kp isolates sequenced in the OH-DART project originated from produce bought from two neighbouring fresh markets (n=205; 34.7%), local chicken, duck, and fish farms (n=272; 46%), community carriage (n=70; 11.8%), hospital inpatients (n=24; 4.1%), and watercourses plus other environmental sources (n=20; 3.4%). 77/591 (13%) of the Kp isolates harboured *iuc*3, 70 of which were recovered from the two fresh markets. The 128 *iuc*3 isolates present in the combined SpARK and OH-DART data are associated with 47 STs (Supplementary Tables 2, 3) representing the breadth of the Kp tree (Supplementary Figure 1).

We generated long-read data and plasmid assemblies for a sub-sample of the *iuc*3 positive isolates in the Italian (n=44, including the single *K. oxytoca* isolate) and Thai collections (n=36), and generated a hybrid assembly for one isolate from a pig in Germany^28^. 79/81 of these plasmid genomes were circularised (Supplementary Table 2). A further 58 assemblies of plasmids containing *iuc*3 were available from public databases, giving a total of 139 plasmid genomes (Supplementary Tables 1, 2). Incomplete assemblies from the SpARK and OH-DART projects and public databases were available for an additional 378 *iuc*3 isolates (Supplementary Tables 1, 3). Whilst the provenance of many of the *iuc*3 isolates in the public databases is unknown, at least 84/517 (16%) from the whole dataset are associated with human infection. *iuc*3 Kp isolates have also been recovered from food products, wastewater^37^, companion animals^26^, and wild animals, including wild boar^28^, invertebrates^38^, white-lipped deer and yak^39^, wild birds, sea lions, and rabbit^40^.

The lineage most commonly associated with *iuc*3 is ST35 (41/517 genomes; 7.9%). This is not simply a consequence of ST35 being very common. Analysis of 29,703 curated Kp genomes using Pathogenwatch^41^ revealed that ST35 represents only 0.87% of all isolates (n=258) and is ranked 17th in terms of frequency.

### Characterising the *iuc*3 plasmids

We first compared the 80 closed plasmids generated for this study from the Italian and Thai studies. It is important to note that differences between these datasets are difficult to interpret as the Thai isolates were selected for resistance to 3^rd^ generation cephalosporins, whilst the Italian isolates were only selected for amoxicillin resistance. Four representative plasmids from the SpARK data and four from the OH-DART data are shown in Supplementary Figure 3, along with plasmid alignments. All of these 80 *iuc*3 plasmids had more than one replicon type, the most common types being IncFIB(K) and IncFII(pKP91) (Supplementary Table 2). These 80 *iuc*3 plasmids ranged in size from 132,430-bp to 365,580-bp. *iuc*3 plasmids from Thailand were larger (mean = 197,626-bp) than the *iuc*3 plasmids from Italy (mean = 159,552-bp) (p<0.001 Wilcoxon Rank-Sum Test) (Supplementary Figure 4A). Ten of the 80 Kp plasmids (12.5%) were >200-Kb, all of which were from Thailand.

ARGs (identified using ABRicate; Supplementary Methods) were more common in the *iuc*3 plasmids from the Thai isolates than those from the Italian isolates (Supplementary Table 2), although, as noted, this can be explained by differences in isolate selection. Ten *iuc*3 plasmids contained at least six ARGs, and nine of these were large Thai plasmids (>200Kb). None of the Italian or Thai *iuc*3 plasmids carried carbapenemase genes, but 7/36 (19.4%) of the Thai plasmids harboured genes encoding extended-spectrum beta-lactamases (ESBLs): *bla*_CTX-M-3_ (n=1), *bla*_CTX-M-55_ (n=1), *bla*_CTX-M-27_ (n=1), and *bla*_SHV-12_ (n=4) (Supplementary Table 2). These seven plasmids meet the definition of ‘convergence’ proposed by Lam et al^10^ based on a Kleborate virulence score of at least 3 (conferred by the presence of *iuc*3), and resistance score of at least 1 (the presence of an ESBL gene). Although originally intended for whole isolates, this definition is equally applicable for single plasmids. The mean size of the seven convergent Thai plasmids was 251,297-bp (range 180,698 - 365,580 bp), which is significantly larger than the non-convergent Thai *iuc*3 plasmids (mean=184,670 bp; range 110,375-287,202 bp) (Wilcoxon Rank-Sum Test, p = 0.00225) (Supplementary Figure 4B). None of the *iuc*3 plasmids in the Italian data were classified as convergent, which again can be partly explained by the difference in isolate selection between the studies.

Of the seven convergent Thai plasmids, six were carried by strains isolated from one of the two fresh markets, and one from an inpatient at the local hospital (Figure 1). All the convergent plasmids were IncFIB(K)/IncFII(pKP91), but the largest plasmid, OH-DART_30005-KC1_2 (365,580-bp), also contained an IncFIA(HI1) replicon. The large size of the convergent plasmids is consistent with them having emerged through hybridisation between co-circulating AMR and *iuc*3 plasmids. To explore this, we identified putative parental plasmids by interrogating all closed plasmid sequences within the dataset based on multiple criteria; ARG profiles, replicon type profiles, AbST, host ST, and nucleotide identity based on BLASTn (see Methods). The majority of the putative parental plasmids showed a high level of nucleotide identity (>98%) to significant fractions of the convergent plasmids, spanning the *iuc*3 and ARG regions. Broadly, putative parental plasmids harbouring *iuc*3 were associated with IncFIB(K), whilst parental AMR plasmids typically carried IncFII(pKP91) replicons and convergent plasmids often carry both.

**Figure 1.**
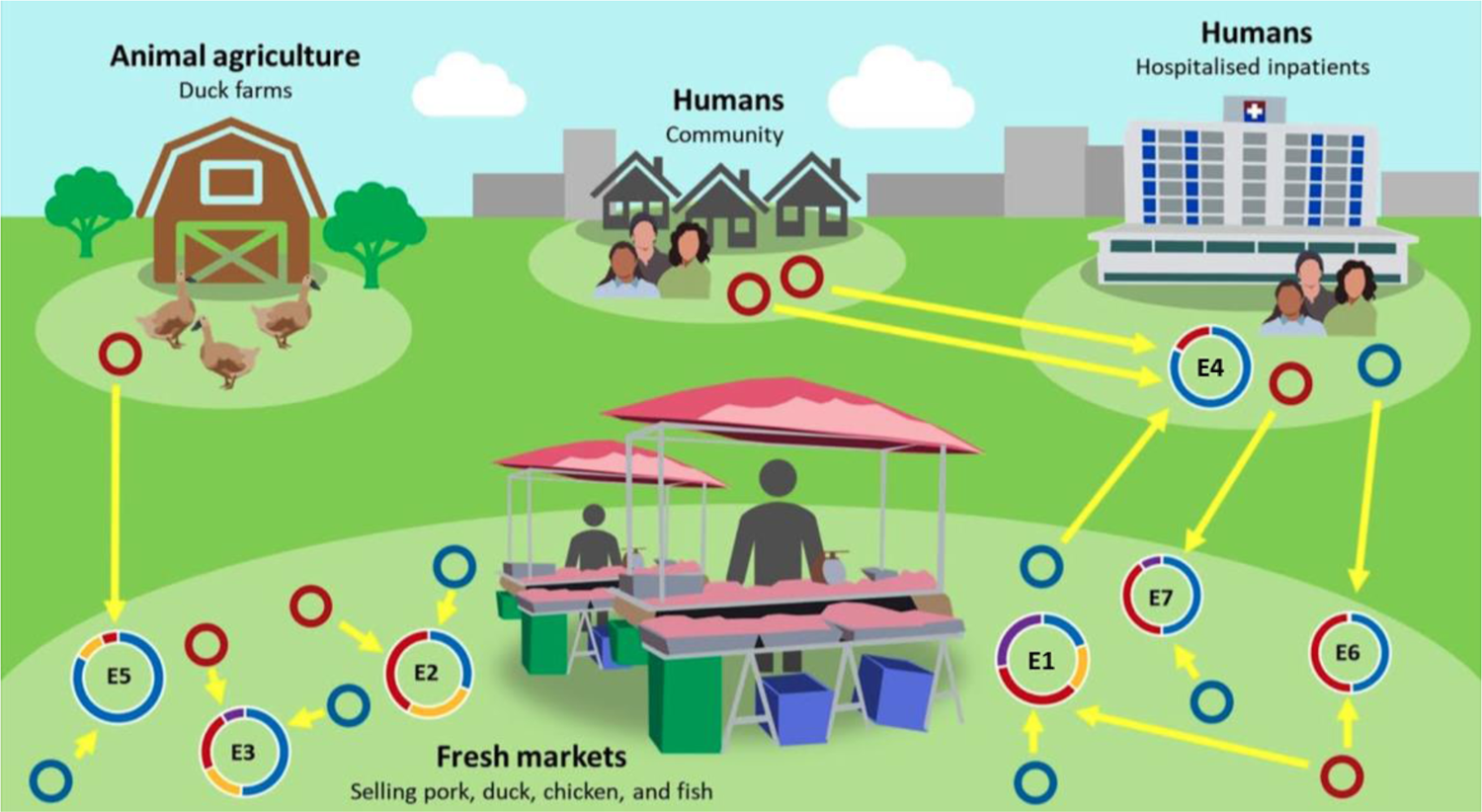
Summary of the seven plasmid convergence events, showing the provenance of the hybrid plasmids and of the putative parental *iuc*3 and AMR plasmids.

A summary of the convergent plasmids and the putative parental plasmids (Events 1-7) is given in Figure 1, and details are provided in Figure 2, Supplementary Figure 5, and Supplementary Table 4. As for the convergent plasmids, the putative parental plasmids were typically carried by isolates recovered from the two neighbouring Thai markets. For example, both putative parental plasmids and the convergent hybrid represented by Event 1 were isolated from the two markets (Figure 2A), and this was also the case for Events 6 and 7 (Supplementary Figure 5C, D). In Event 4 (Supplementary Figure 5A), two different AMR plasmids from community carriage were identified as potential donors, one of which was harboured by an *E. coli* isolate that was also sequenced as part of the OH-DART project.

**Figure 2A (Event 1).**
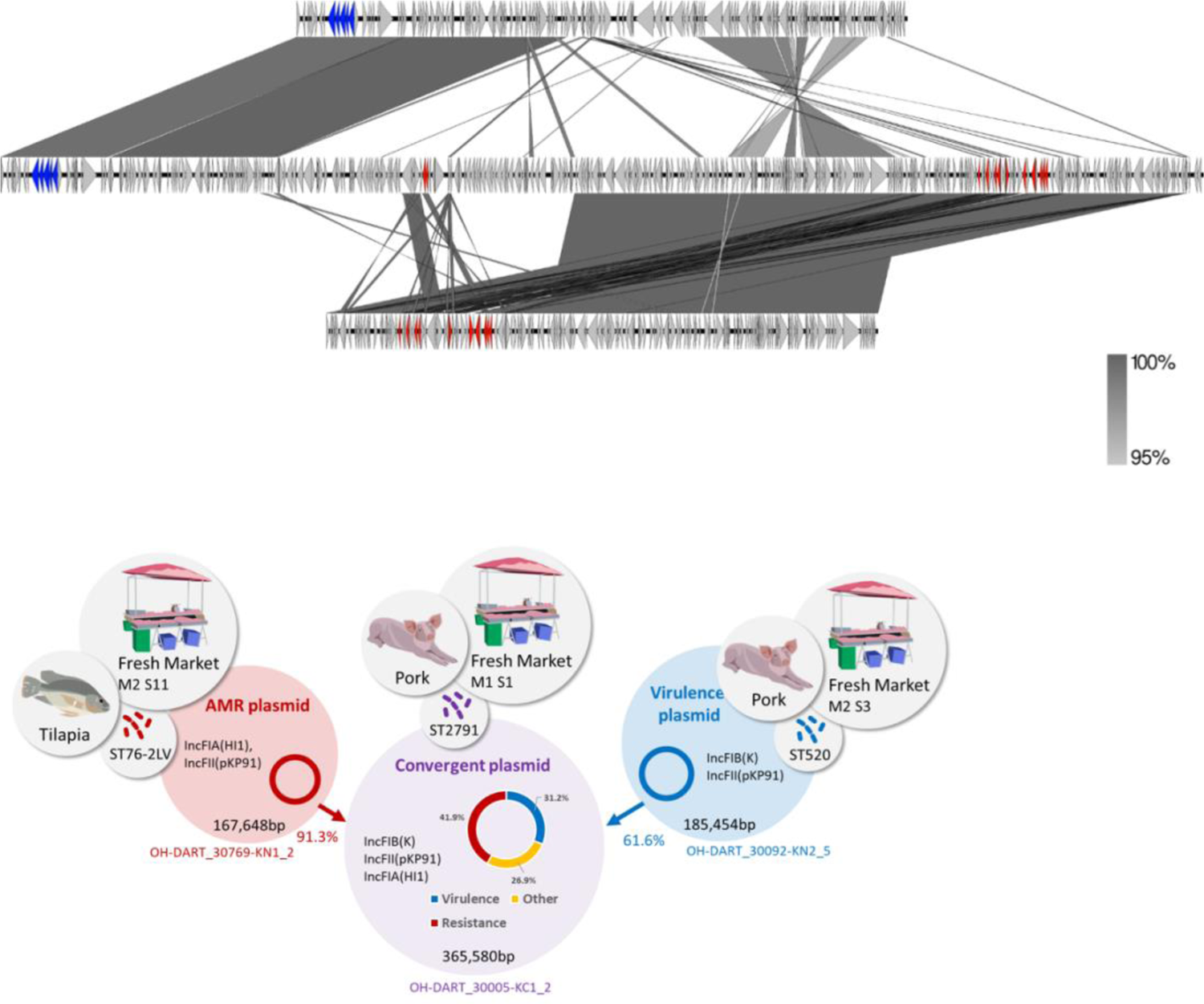
The convergent plasmid OH-DART_30005-KC1_2 (middle) is the largest fully sequenced *iuc*3 plasmid in the dataset (365580-bp) and was used as a reference for mapping and SNP calling. A circular representation of this plasmid showing other genome features is provided in Supplementary Figure 2A. This plasmid has replicon types IncFIB(K), IncFII(pKP91), and IncFIA(HI1), and was isolated from an ST2791 isolate recovered from pork meat. 31.2% of this plasmid showed high level of nucleotide identity (>98%) to 61.5% of the *iuc*3 plasmid OH-DART_30092-KN2_5 (ST520; IncFIB(K), IncFII(pKP91)) which was recovered from pork meat (top). 41.9% of the large convergent plasmid showed >98% sequence identity to 91.3% of the AMR plasmid OH-DART_30769-KN1_2 (bottom) (ST76-2LV; IncFIA(HI1), IncFII(pKP91)). The ARG profile of the AMR plasmid OH-DART_30769-KN1_2 is *aac(3)-IId*^; *aadA17*\*; *strA*.v1; *strA*.v1^; *strB*.v1; *strB*.v1 *qnrS1*; *erm*(42)*; *lnuF*.v1; *sul2*; *tet*(A).v2; *bla*_SHV-12_. The convergent plasmid also harbours the same ARG profile, but with the addition of an *erm* gene. We note that 26.9% of the convergent plasmid does not correspond to either of the putative parental plasmids, thus material from at least one other plasmid has contributed to this large genome.

**Figure 2B (Event 2).**
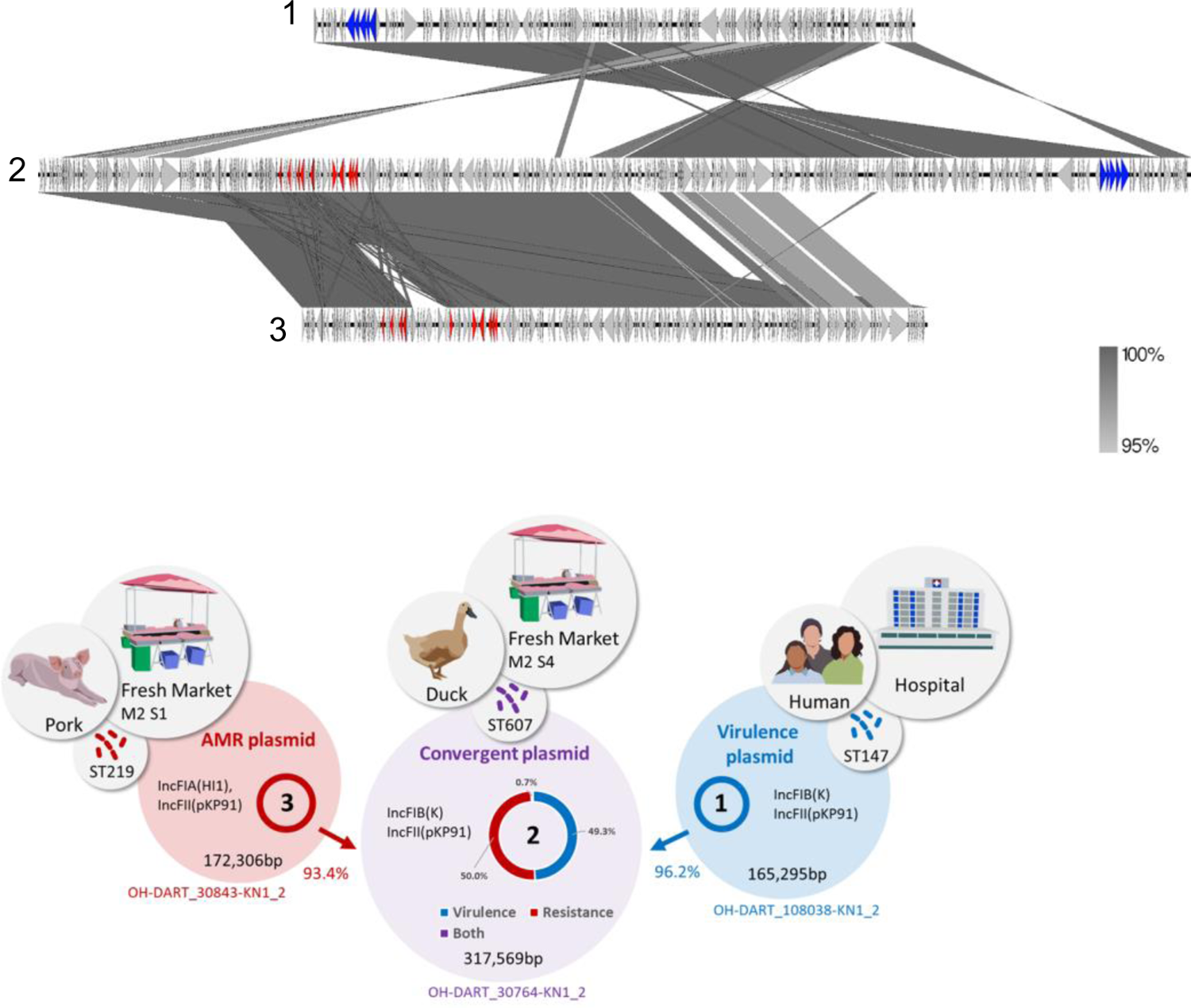
This example illustrates the hybridisation of an AMR plasmid from the market (OH-DART_30843-KN1_2; ST219) with an *iuc*3 plasmid from the faecal flora of a hospital inpatient (OH-DART_108038-KN_2). The clinical isolate harbouring the *iuc3* plasmid corresponds to the high-risk clone ST147 (Rodrigues et al 2022 DOI: 10.1099/mgen.0.000737), although this isolate does not contain significant ARGs. The convergent hybrid plasmid, OH-DART_30764-KN1_2 (ST607), which was isolated from duck meat from the fresh market, is an almost perfect hybrid of these two plasmids. 93.4% of plasmid OH-DART_30843-KN1_2 and 96.1% of plasmid OH-DART_108038-KN_2 each share >99% nucleotide identity with 50% of OH-DART_30764-KN1_2 plasmid. OH-DART_30843-KN1_2 and the convergent plasmid OH-DART_30764-KN1_2 share an identical ARG profile: *aac(3)-IId*^; *aadA17*\*; *strA*.v1^; *strB*.v1; *qnrS1*; *lnuF*.v1; *floR*.v2*; *sul2*; *tet*(A).v2; *bla*_SHV-12_. The arrows refer to plasmid hybridisation, and do not denote transmission pathways.

**Figure 2C (Event 3).**
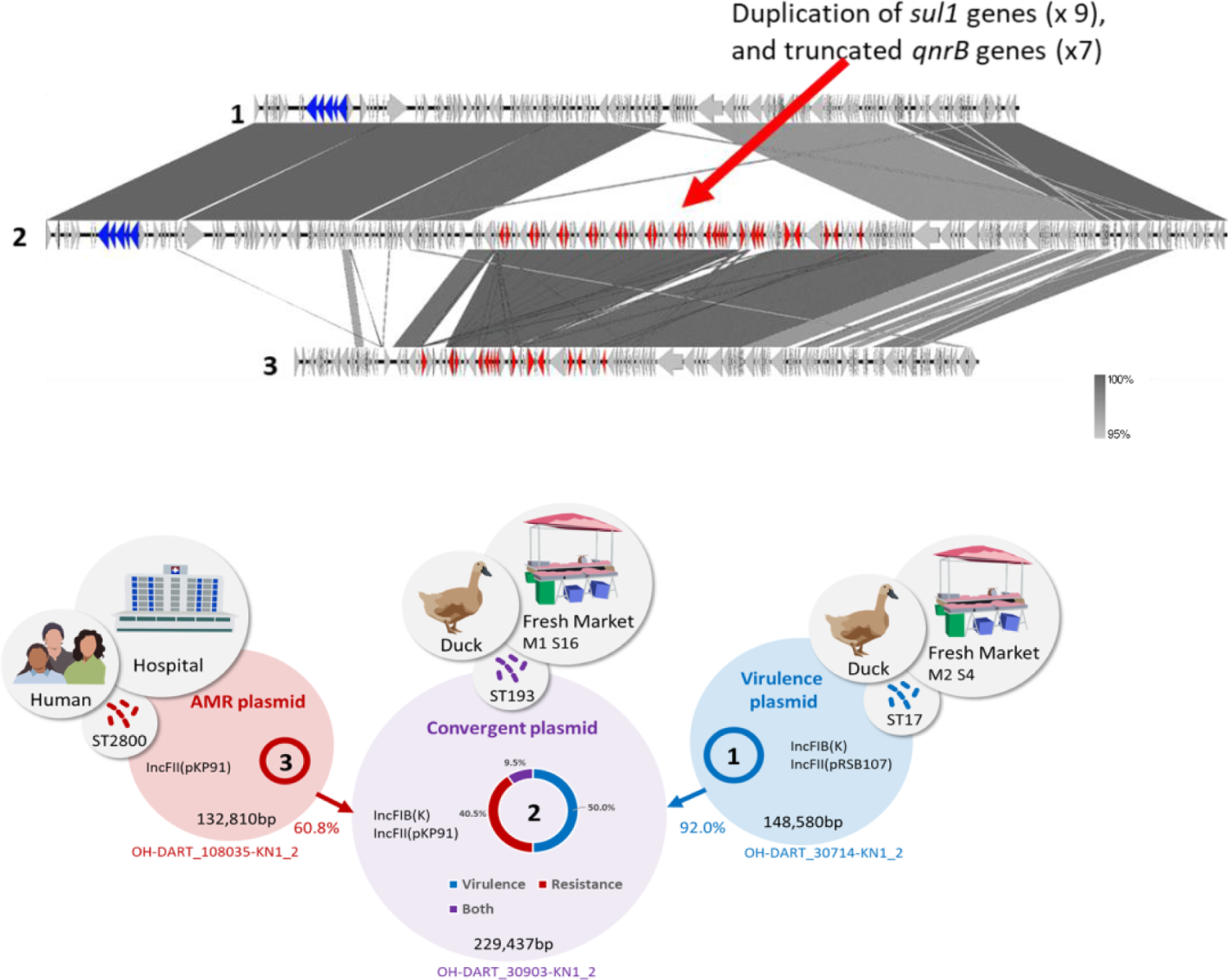
In this case the putative parental AMR plasmid originates from a stool sample from a hospital inpatient whilst the convergent and putative parental *iuc*3 plasmids were carried by isolates from the markets. 59.5% of the convergent plasmid OH-DART_30903-KN1_2 (ST193) shares >98% nucleotide identity with 91.9% of the *iuc*3 plasmid OH-DART_30714-KN1_2 (ST17). 50% of plasmid OH-DART_30903-KN1_2 shares >98% identity with 60.7% of plasmid OH-DART_108035-KN1_2 (ST2800), and these latter two plasmids share identical ARG profiles: *aac(6’)-Ib-cr*.v2; *aadA16*; *aph3-Ia*.v1; *qnrS1*; *mphA*; *floR*.v1; *arr-3*; *sul1*; *tet*(A).v2; *dfrA27*; *bla*_TEM-1D_.v1; *bla*_CTX-M-3_; *qnrB20*\*-10%. However, curiously, the *sul*1 and truncated *qnr*B20 genes have been duplicated in the convergent plasmid, resulting in 9 tandem copies of the former, and 7 tandem copies of the latter. The arrows refer to plasmid hybridisation, and do not denote transmission pathways.

**Figure 2D.**
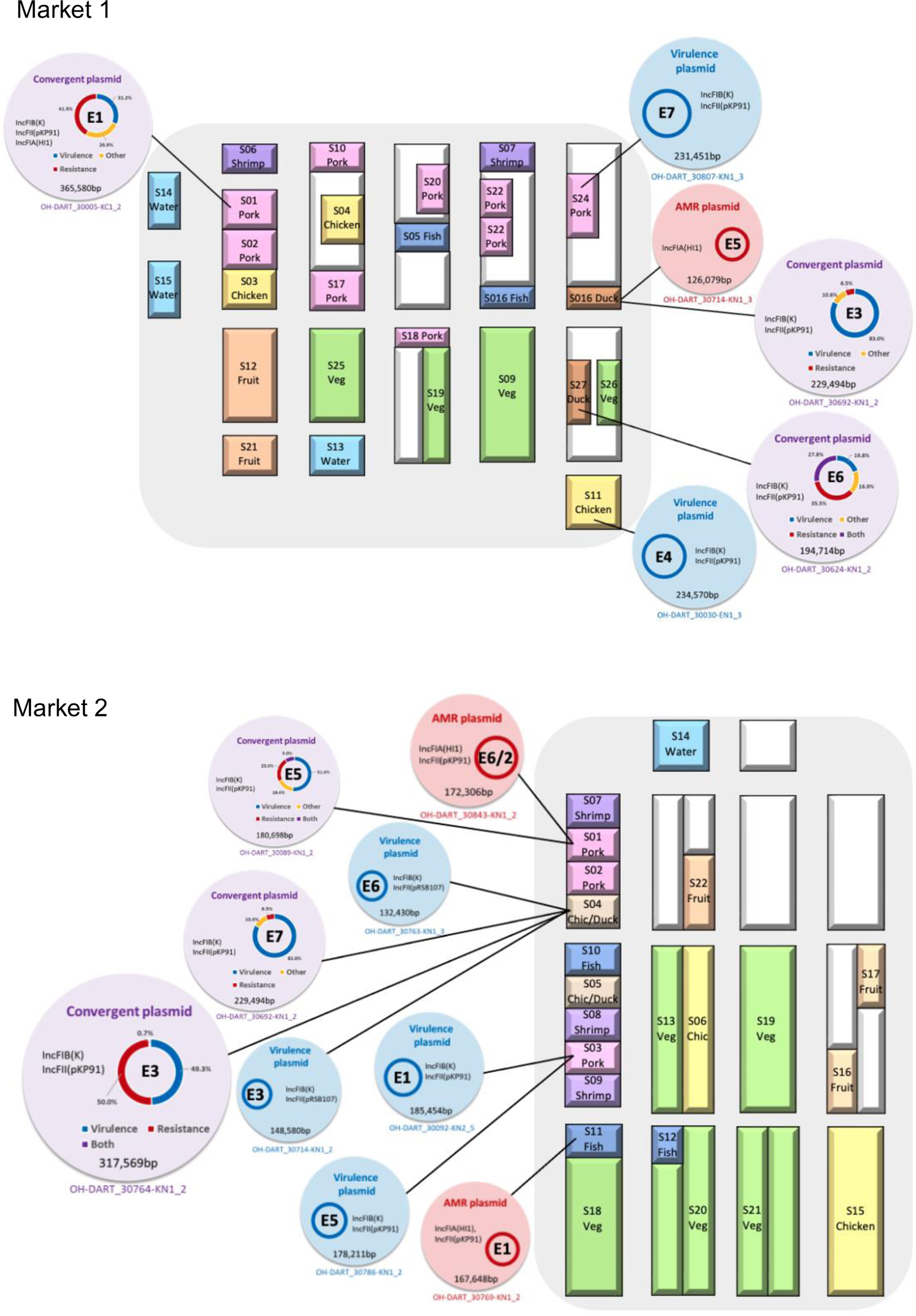
Locations of the sources of convergent plasmids and the putative parental plasmids on the two Thai markets. The Event (‘E’) numbers are shown within each plasmid. Red represents the contribution from the AMR plasmid, blue from the *iuc*3 plasmid. OH-DART_30843-KN_2 is the putative AMR parental plasmid in Events 2 and 6. Events 4-7 are shown in Supplementary Figure 5.

Analysis by MOB-typer revealed that all convergent plasmids are predicted to be conjugative, as are all putative parental plasmids, with the exception of the AMR plasmid from *E. coli* (mobilizable) and a small resistance plasmid from a duck farm isolate (non-mobilizable) (Supplementary Table 4). Events 2, 3 and 4 indicate a flow of plasmid material between isolates from the local market, community carriage and clinical isolates recovered from inpatients at the nearby hospital (Figures 1 and 2, Supplementary Figure 5, Supplementary Table 4).

There are likely to be many other plasmid hybridisation events in the data that do not meet the definition of convergence as they do not carry an ESBL or carbapenemase gene. One such example is shown in Supplementary Figure 6, where a single ST7513-1LV isolate carries both an *iuc*3 plasmid and second resistance plasmid harbouring multiple ARGs including *bla*_CMY-2_. Although *bla*_CMY-2_ confers resistance to 3^rd^ generation cephalosporins, it is not commonly encountered in clinical settings and is not assigned as an ESBL gene. A second isolate of the ST7513-1LV clone harbours a near-perfect, and very large (287,202 bp), hybrid of these two plasmids, including all ARGs from one parent, the *iuc*3 locus from other and all 5 replicon types found in the parental plasmids (Supplementary Figure 6).

To further explore the global diversity of *iuc*3 plasmids, we combined the 80 *iuc*3 complete plasmid sequences from Italy and Thailand, and the single plasmid from Germany^28^, with an additional 58 from the public domain. Two of the 58 public *iuc*3 plasmid sequences are convergent: pKPC-063001 (359,625 bp; accession: MZ156798.1) harboured by a clinical ST11 isolate from China and contains a *bla*_KPC-2_ gene^42^, and pKPT877-hybrid (293,653 bp; accession: CP084242.1) which has a *bla*_OXA-181_ carbapenemase gene and was harboured by a clinical ST43 isolate from China^43^. We carried out a gene content analysis of all 139 closed plasmid genomes using Roary (Supplementary Methods). The total number of genes was 1877, of which 26 ‘backbone genes’ were present in at least 136/139 (>97.5%) of the plasmids (Supplementary Figure 7). We note a high frequency of insertion sequences among the 139 plasmids (Supplementary Table 5). The most common IS family is IS*3*, of which there are 562 copies, and other common families are IS*110* (n=297), IS*481* (n=257) and IS*6* (n=180). The majority of the IS*6* family are likely to be IS*26* which are known to play a key role in the transfer and insertion of resistance gene clusters.

### Clustering and phylogenetic analysis

We used mge-cluster to compare the 139 *iuc*3 plasmids in the context of nearly 3000 *iuc*3 negative plasmids from the SpARK and OH-DART data sets (Supplementary Figure 8). Half (70/139) of the *iuc*3 plasmids correspond to two neighbouring clusters (#9; n=35 and #25; n=35), with the others mostly corresponding to a looser ‘cluster of clusters’ (#s 23, 27, 28), or not assigned to a cluster (#-1, n=23). The *iuc*3 locus itself does not appear to be highly mobile, as only a small number of *iuc*3 plasmids are dispersed among major clusters. Exceptions include three convergent plasmids (OH-DART_30005-KC1_2, OH-DART_30764-KN1_2, and OH-DART_30624-KN1_2) that correspond to the large cluster #49, but this is likely to be the result of plasmid hybridisation rather than mobility of the *iuc*3 locus (Figures 2A,B, Supplementary Figure 5C; Events 1, 2 and 6). From the public data, plasmid SWHEFF_62, which was isolated from wastewater in Hong Kong^37^, belongs to the large cluster #38, and KP_NORM_BLD_2015_115359_unnamed_1, a clinical isolate from Norway, belongs to cluster #60.

Of the 35 *iuc*3 plasmids assigned to cluster #9, eight are from the Thai dataset, with the remaining 27 from the public domain. All are of Asian origin, being sampled from China (n=25), Thailand (n=8), Hong Kong (n=1) and Vietnam (n=1). Cluster #9 contains 4 convergent plasmids, two from the Thai study, and two from the public data. 16/39 (45.7%) of the cluster #9 *iuc*3 plasmids are from humans, with seven known to be from clinical isolates. Although none of the *iuc*3 plasmids from the Italian study cluster in this group two *iuc*3-negative plasmids from this study do, both of which were harboured by clinical isolates: SPARK_356_C1_2 (which also harbours a *bla*_CTX-M-15_ gene), and SPARK_551_C1_4^26^. In contrast to cluster #9, the 35 *iuc*3 plasmids in cluster #25 are from diverse geographical sources (Thailand, n=12; Italy, n=10; Norway, n=6; China, n=3; Laos, n=3; USA, n=1), and 9/35 (25.7%) are associated with humans, mostly from clinical isolates. The *iuc*3 plasmid from the *K. oxytoca* isolate clusters in this group and is not diverged from the Kp plasmids, indicating recent inter-species transfer.

To check the robustness of the clusters, we mapped the 139 plasmid genomes to the reference plasmid, and built a tree based on SNPs (Supplementary Methods; Supplementary Figure 9). There are only minor inconsistencies between clusters and the resulting phylogeny. Clusters #25 and #9 are clearly resolved on the tree (assigned Groups 2 and 3 respectively). Group 3 (cluster #9) plasmids are the most similar to each other and contain more ARGs than other plasmids (Supplementary Figure 10A). The overlapping clusters #23, #27 and #28, and those not assigned to a cluster (−1), form a more diverse third group (assigned Group 1), represented by multiple replicon types (Supplementary Figure 10B). This tree is also consistent with a simplified AbST scheme (Supplementary Methods), with Group 3 corresponding to AbST 23, Group 2 to AbST 25 and Group 1 to AbSTs 43 and 86 (Supplementary Figures 10C, 11A,B, Supplementary Table 6).

Differences in gene content between the three groups are shown Supplementary Figure 12, and genes that are enriched within Group 3, which are associated with clinical Asian isolates, are given in Supplementary Table 7. These include genes associated with Type-4 secretion system (T4SS) (*trbC*), gene regulation (e.g. *cspA*, *repA*), Type 1 R-M system, metalloproteases (*sprT*), IS*1* and IS*3* family transposases, a complete *fec* operon associated with iron uptake, a histidine degradation pathway (*hut*), and an *ars* operon encoding resistance to arsenic. This latter operon includes a transcriptional repressor, *arsR,* that has a global regulatory role beyond the *ars* operon^44^.

Finally, we mapped short reads from all 517 *iuc*3 isolates (including those for which plasmid assemblies are not available) to the reference plasmid OH-DART_30005-KC1_2 (the largest *iuc*3 plasmid in the data) and built a SNP-based tree (Figure 3a). The tree and metadata are available to inspect in a Microreact project at https://microreact.org/project/gibbonetal-517-iuc3. The three main plasmid groups are also evident on this tree, and the AbST data also remains consistent. For example, almost all Group 3 plasmids are in the simplified AbST23 group except for four likely artefactual exceptions (Supplementary Note). A breakdown of each group by geographical and ecological source is given in Supplementary Table 8, and this further supports the view that the Group 3 plasmids are associated with Asian isolates, as 140/158 (88.6%) of the plasmids in this group are from Asian origin (Supplementary Table 8; Figure 3b). There are only 11 plasmids of European origin clustering in Group 3, and ten of these are associated with humans, and at least eight of them from clinical isolates (Supplementary Figure 13). The earliest example of a Group 3 plasmid isolated in Europe was in 2012 in the UK; these plasmids have therefore been occasionally imported into Europe from Asia for at least 10 years.

**Figure 3.**
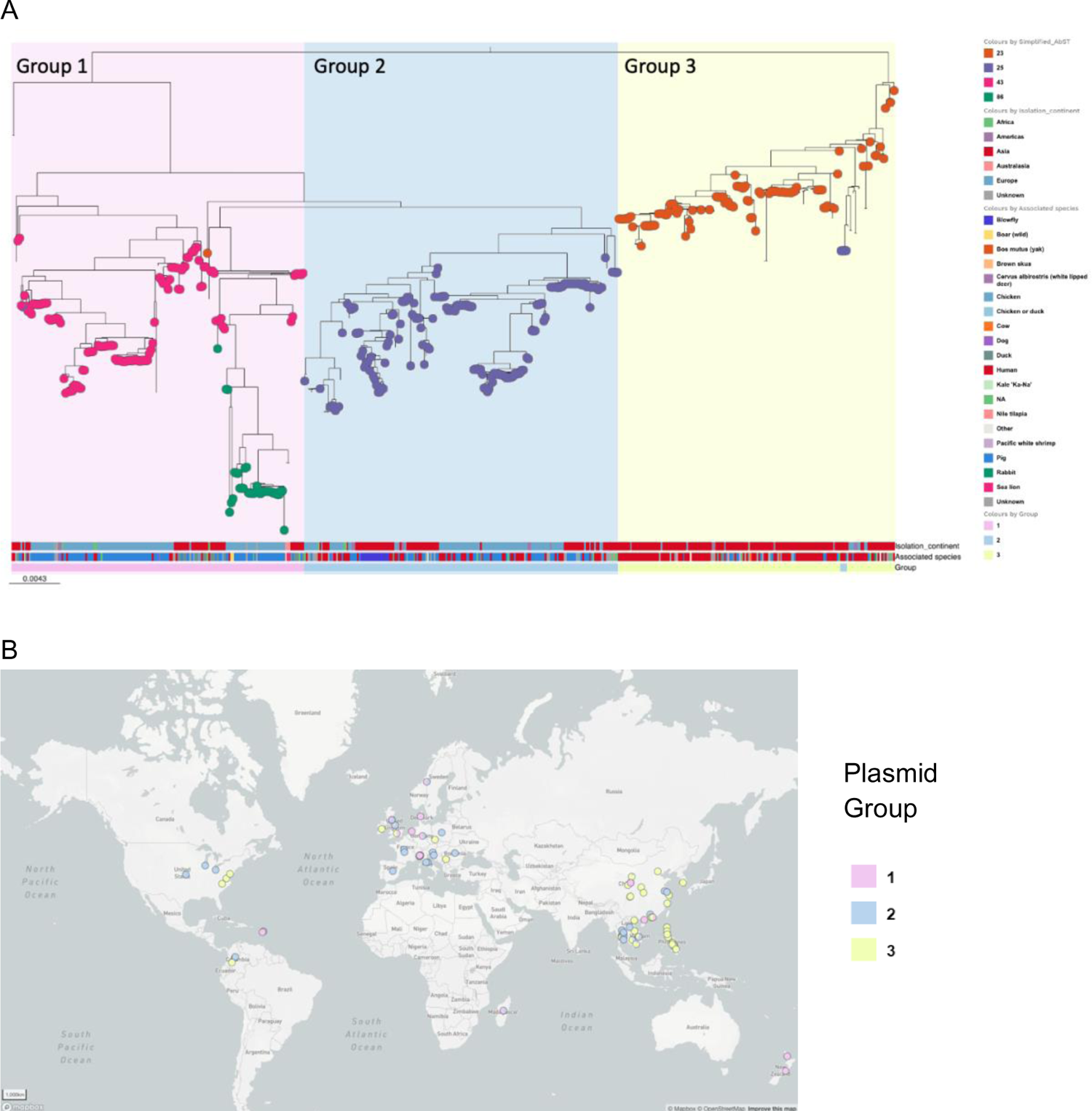
Phylogeographical analysis of 517 *iuc3* plasmids. **A.** SNP tree of 517 genomes coloured by simplified AbST, showing continent of isolation and associated species. **B.** The geographical distribution of the three groups of *iuc*3 plasmids. Group 3 plasmids (yellow) are predominant in China, the Philippines and Southeast Asia. Four AbST25 plasmids pMR0617aac, GCF_002752955.1, 480738_p and OH-DART_30092-KN2 are assigned as Group 2 due to their position on the tree of 139 plasmids (Supplementary Figures 9, 10c).

Overall, 66% of the Group 3 plasmids were associated with human isolates (104/158), and only 5% with animal isolates (8/158). In contrast Group 1 is associated with animal isolates from Europe (9% human, 65% animal,66%; Europe, 26% Asia). Group 2 is not strongly associated with either continent or source (35% human, 40% animal; 51% Europe, 43% Asia). The bulk of the *iuc*3 Thai plasmids from the OH-DART study were isolated from markets and were assigned as ‘foodstuffs’. These are scattered across the three groups (Supplementary Figure 14).

Finally, we checked the distribution of STs across the larger global tree of 517 isolates (Supplementary Figure 15A). As noted earlier, 41 of the plasmids were harboured by ST35 Kp hosts (or single locus variants), more than any other ST. The *iuc*3 plasmids carried by ST35 isolates represent all three groups and are recovered from both clinical and animal isolates from both Europe and Asia. We compared the tree of just the 41 ST35 plasmids with the corresponding chromosomal tree in order to examine the degree of co-diversification between host and plasmid (Supplementary Figure 1B,C). This revealed a sub-lineage of 16 closely related *iuc*3 ST35 isolates, 15 from Thailand and one from China, all of which harbour Group 3 plasmids. Thus, an Asian sub-variant of ST35 acquired a Group 3 *iuc*3 plasmid and this has been stably maintained in this lineage and disseminated throughout Asia. There are no consistencies between the plasmid and chromosomal trees when only plasmids outside of Group 3 are considered.

## Discussion

The rapid and highly reticulate evolution of plasmids poses serious challenges for clustering, typing or phylogenetic reconstruction^45^. By focusing on a set of plasmids selected from our sequence data simply on the basis of the presence of a single virulence locus, *iuc*3, we have surreptitiously struck upon a ‘Goldilocks’ level of diversity, whereby there is enough variation to identify robust groups, but not so much that the phylogenetic signal is mostly conflicting. The consistency between clustering and phylogenetic analysis bodes well for future studies on plasmid epidemiology, and the resolved clusters are mostly consistent with other plasmid features, and in particular AbST which indicates stable co-diversification between the plasmid and the *iuc*3 locus. The plasmid clusters we defined are therefore both evolutionary meaningful and epidemiologically useful. The oldest plasmid in our data was associated with a Danish isolate sampled in 1952, meaning that the association between *iuc*3 and these plasmids is over 70 years old.

Analysis of plasmid diversity enabled us to confidently delineate a distinct *iuc*3 plasmid sub-variant (Group 3/Cluster 9) that is associated with clinical isolates circulating throughout Asia. It is unclear to what extent the association between Group 3 plasmids and clinical isolates is driven by the presence of the *iuc*3 locus itself, or other genes specifically associated with this plasmid group. Group 3 plasmids are associated with the *iuc*3 sub-variant AbST23, which is not notably diverged from other *iuc*3 AbST types making it less likely that it enhances virulence compared to other sub-variants. In addition to multiple ARGs (including in 2 cases carbapenemase genes), these plasmids also contain global regulators, and operons for iron uptake and arsenic resistance. The presence of two Italian *iuc*3-negative plasmids from clinical isolates that also cluster with this group further points to a role for other genes on the plasmid. The ability to generate a meaningful plasmid tree sheds light on other aspects of the data. For example, the *iuc*3 plasmids from the Thai markets (2 Km apart) represent the full global diversity of these plasmids. This remarkable observation suggests that *iuc*3 plasmids have been circulating in this region for a long time, presumably in pigs or other animals. Although *iuc*3 is found in a broad range of STs representing the breadth of the Kp phylogeny, indicating frequent plasmid transfer, we do note an association between *iuc*3 and ST35 isolates that cannot be explained by simple clonal spread. The diversity of *iuc*3 plasmids within ST35 strains, and the fact that *iuc*3 carrying ST35 isolates are found globally and from diverse sources, indicates multiple plasmid acquisitions within this lineage. However, we also note a sub-lineage of Asian ST35 isolates that has acquired - and become associated - with Group 3 *iuc*3 plasmids. From the relatively low level of diversity of this sub-variant (both of the chromosome and of the plasmid) this association probably arose recently, and we note that the oldest example in the data is from 2015.

Of the 36 fully sequenced *iuc*3 plasmids in the Thai study, 7 contained an ESBL gene and so met the definition of ‘convergent’. Six of these plasmids were harboured by isolates recovered from the market and emerged through hybridisation with AMR plasmids also mostly recovered from the markets. All seven hybrid plasmids are predicted to be conjugative, thus can act as vehicles for the simultaneous transmission of both virulence and resistance traits. Whilst convergence through plasmid hybridisation is a well-known phenomenon for Kp^39^ to our knowledge this is the first time that multiple events have been described outside of a clinical setting, and the first time that population-scale data have been used to identify putative parental plasmids.

Frequent plasmid hybridisation also challenges our view of what constitutes ‘transmission’. There were two isolates from hospital patients in the Thai data harbouring *iuc*3 plasmids, from a total of only 20 Kp hospital isolates sampled in the OH-DART study. Potential links between the patient and market isolates based on *iuc*3 plasmids are apparent that would have missed on using standard genomic epidemiology. For example, in event 2, the *iuc*3 plasmid OH-DART_108038-KN1_2 which was isolated from the faecal flora of a hospital inpatient, makes up almost exactly half of the convergent plasmid OH-DART_30764-KN1_2 isolated from duck meat from the market. It is not possible from this to infer direct transmission between the hospital and the market, and if there is an epidemiological link, it is likely that there are unsampled intermediate links in the chain. Nevertheless, this example demonstrates how deep sampling of a defined population, the generation of closed assembled genomes (i.e. long-read data), and careful bespoke analysis can reveal more cryptic and indirect epidemiological connections involving plasmids.

We posit that such plasmid hybridisation events happen very frequently in natural populations, although the fitness consequences of any given event for both the plasmid and the host, and the commensurate public health threat remain unclear. Further questions persist, concerning the underlying mechanisms, and how to predict the likelihood that any given pair of plasmids might hybridise in natural populations. The identification of putative parental plasmids will shed light on this, and in particular the role of insertion sequences, as these are known to be a key driver of dynamics in plasmid genomes. A recent report describing a convergent pLVPK-like plasmid (*bla*_NDM-1_ plus *iuc*/*rmp*A) carried by an ST11 isolate points to mediation via multiple copies of IS*26*-like elements^46^, and this plasmid has been found to be transmitting from hospitals to the aqueous environment^47^. Structural rearrangements in plasmids mediated by IS*26*-like elements have also recently been noted in *Salmonella enterica* Serovar Typhimurium^48^, and these elements are known to play an important role in the dissemination of key ARGs, including *tet*(X) which encodes resistance to tigecycline^49^.

In conclusion, here we use a combination of sequencing and interrogation of public databases to conduct a comprehensive analysis of the diversity and distribution of Kp plasmids harbouring the virulence locus *iuc*3. The use of data from ‘One-Health’ settings provides important context, and we argue that the emergence of virulence, as well as AMR, needs to be considered within this framework. The study provides an example for the utility of closed assemblies for studying the evolutionary and epidemiological dynamics of plasmids on global scales.

## Acknowledgements

We are grateful to Silvia Argimón for help and advice and to Guido Werner and Sébastien Breurec for the provision of unpublished data on the isolates from Germany and the French West Indies. We are also grateful to I-Ting Tu, Ellen Cardwell and Lily Feil for technical assistance.

## Data Availability

Data generated by this project are deposited under Project numbers PRJEB66363 and PRJEB66356. The tree of 571 *iuc*3 plasmids, and associated metadata, is available to explore at https://microreact.org/project/gibbonetal-517-iuc3.

## Supplementary Figures

**Supplementary Figure 1.**
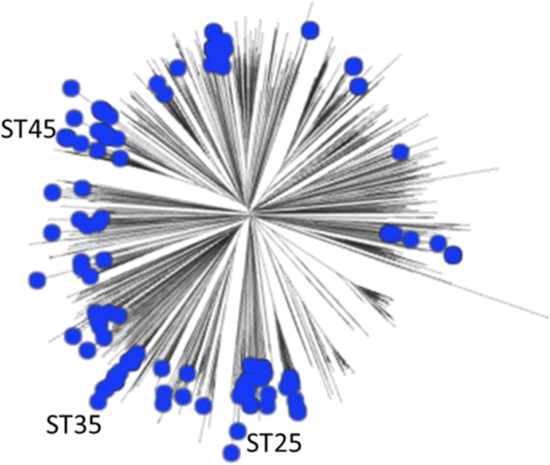
Mashtree of 2296 Kp isolates from Italy and Thailand. The blue nodes indicate the presence of *iuc*3. The three STs most commonly associated with *iuc*3 are indicated.

**Supplementary Figure 2.**
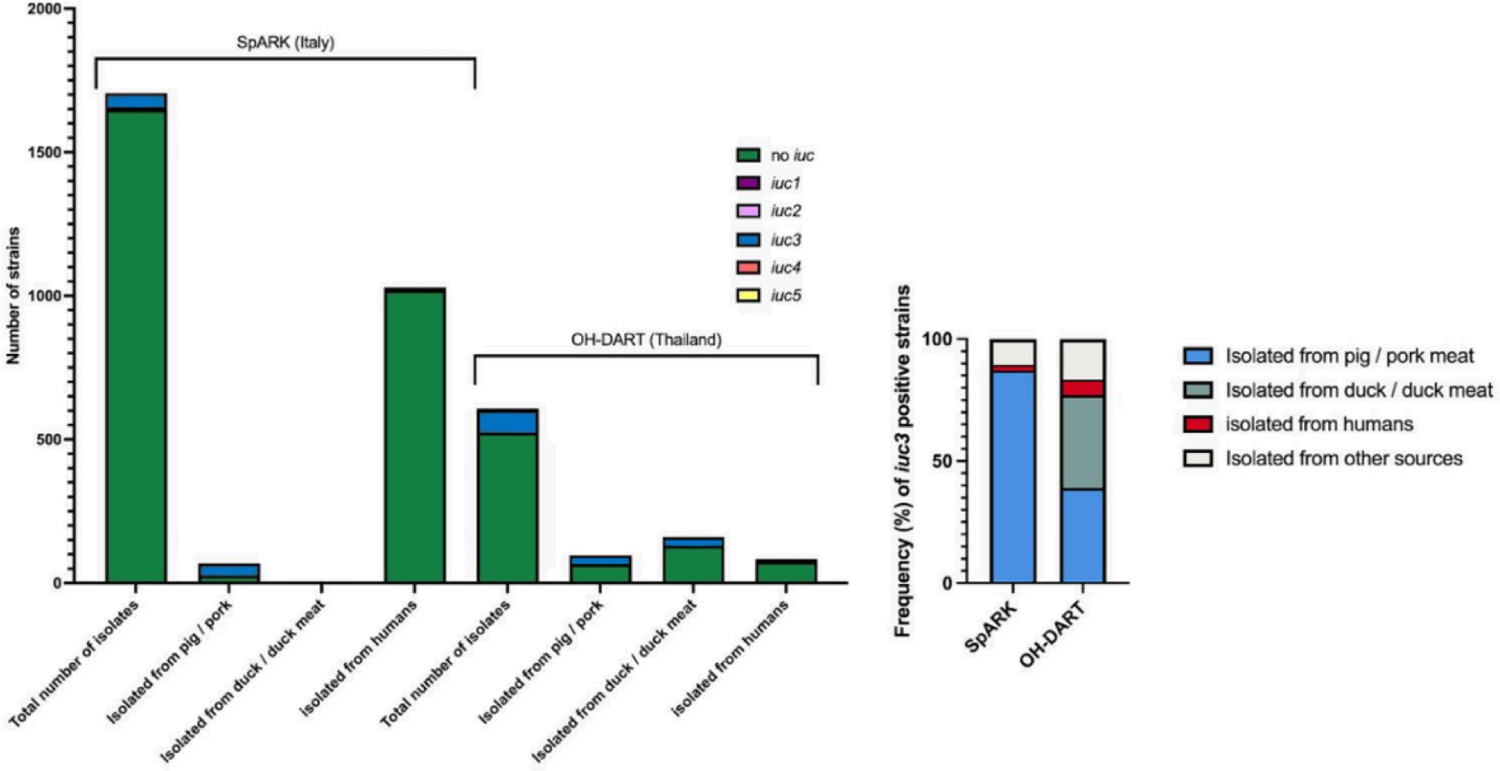
Sources and prevalence of *iuc*3-positive isolates in the Italian (SpARK, n=45) and Thai (OH_DART, n=36) datasets.

**Supplementary Figure 3.**
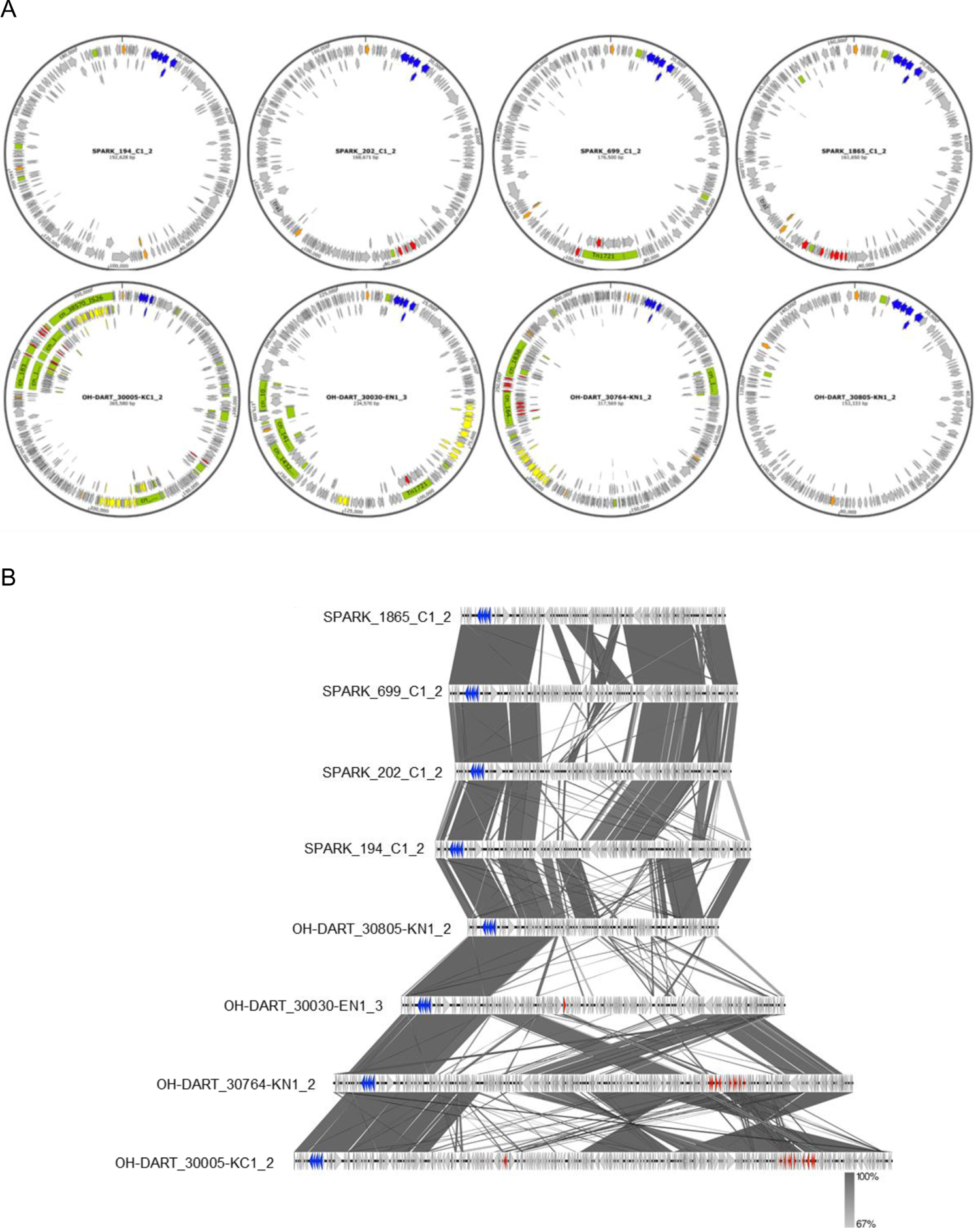
Representative *iuc3*-carrying plasmids from Thailand and Italy. **A.** Circular sequence maps showing the *iuc*3 locus (blue), ARGs (red), *rep* gene (orange), heavy-metal resistance (yellow) and IS/Tn (green). **B.** Alignment of the same plasmids, showing *iuc*3 locus (blue) and ARGs (red).

**Supplementary Figure 4.**
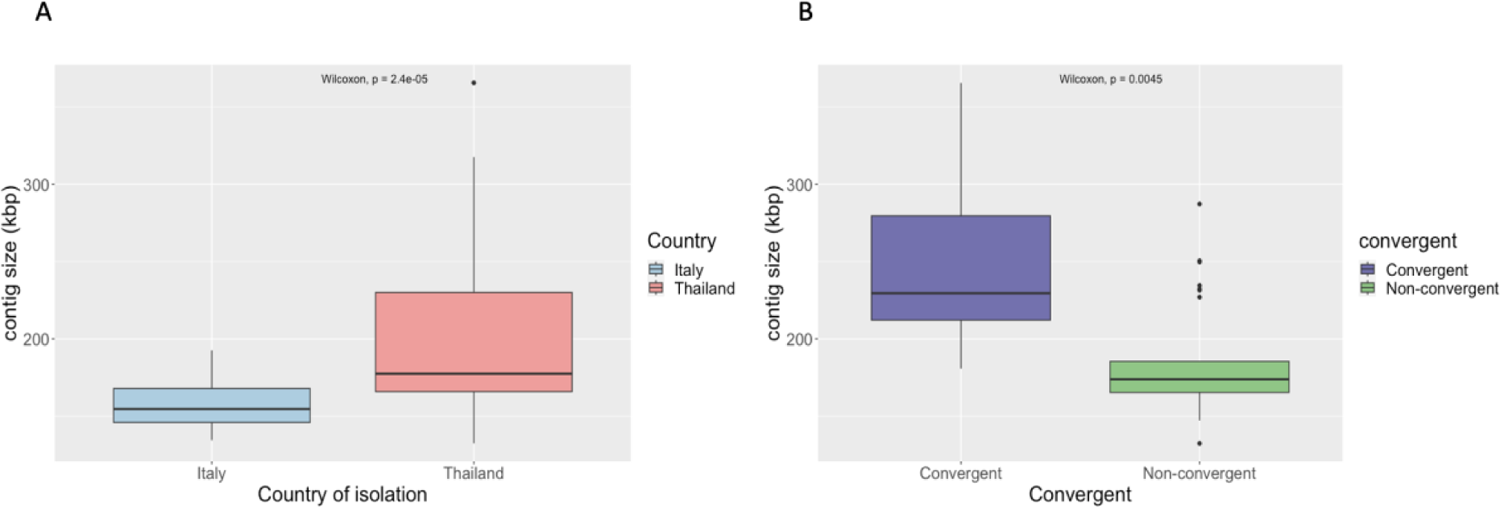
Size comparisons of *iuc3* plasmids **A.** from Italy (N= 44, blue) and Thailand (N= 36, red) and **B.** Thai convergent plasmids (N=7, purple) and Thai non-convergent plasmids (N=29, green). Both comparisons show significant difference by a Wilcoxon Rank-Sum Test (P<0.001).

**Supplementary Figure 5A (Event 4).**
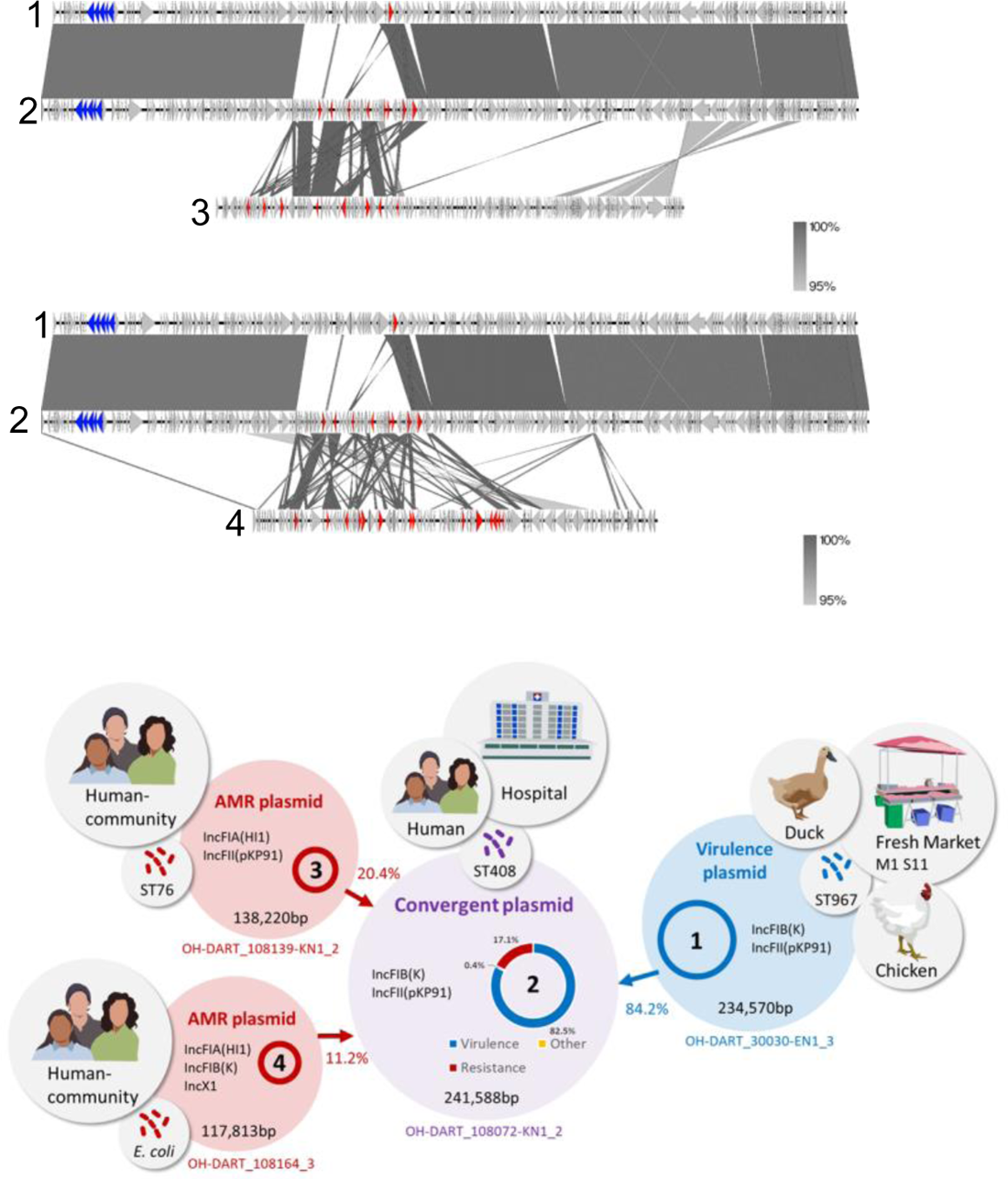
A 241,588-bp convergent *iuc*3 plasmid (OH-DART_108072-KN1_2; ST408) was carried by an isolate recovered from the faecal flora of an inpatient at Ban Leng hospital. This plasmid has the following ARG profile: *aadA2*, *strAB*, *qnrS1*, *sul2*, *tet*(A), *dfrA12*, *bla*_LAP-2_, *bla*_CTX-M-27_. 82.5% of this plasmid showed high homology (>98% identity) to an *iuc*3 plasmid (OH-DART_30030-EN1_3; ST35) isolated from one of the fresh markets, which does not contain any ARGs. The ARG profile of plasmid OH-DART_108139-KN1_2; (ST76), from a community carriage isolate, shows some overlaps with that of the convergent plasmid: *aac(3)-IId*^, *qnrS1*, *mphA*, *floR.v1*\*, *sul2*, *tet*(A).v1; *dfrA14*.v2*; *bla*_LAP-2_; *bla*_SHV-12*_, and 20.4% of this plasmid showed >98% identity to 11.4% of the convergent plasmid. The *bla*_CTX-M-27_ ESBL gene in the convergent plasmid may have been derived from a parent similar to the IncFIA(HI1), IncFIB(K), IncX1 plasmid (OH-DART_108164-EN1_3), which was harboured by an *E. coli* strain also recovered from local community carriage. This plasmid contains a *bla*_CTX-M-14_ gene which is >99% identical to the *bla*_CTX-M-27_ gene on the *E. coli* plasmid. 5.4% of the convergent OH-DART_108072-KN1_2 plasmid shows >98% nucleotide identity to 11.1% plasmid OH-DART_108164-EN1_3. This event points to the potential flow of plasmid material between the fresh market, community carriage, hospital carriage, as well as highlighting the possibility of the transfer of ARGs between *K. pneumoniae* and *E. coli* plasmids.

**Supplementary Figure 5B (Event 5).**
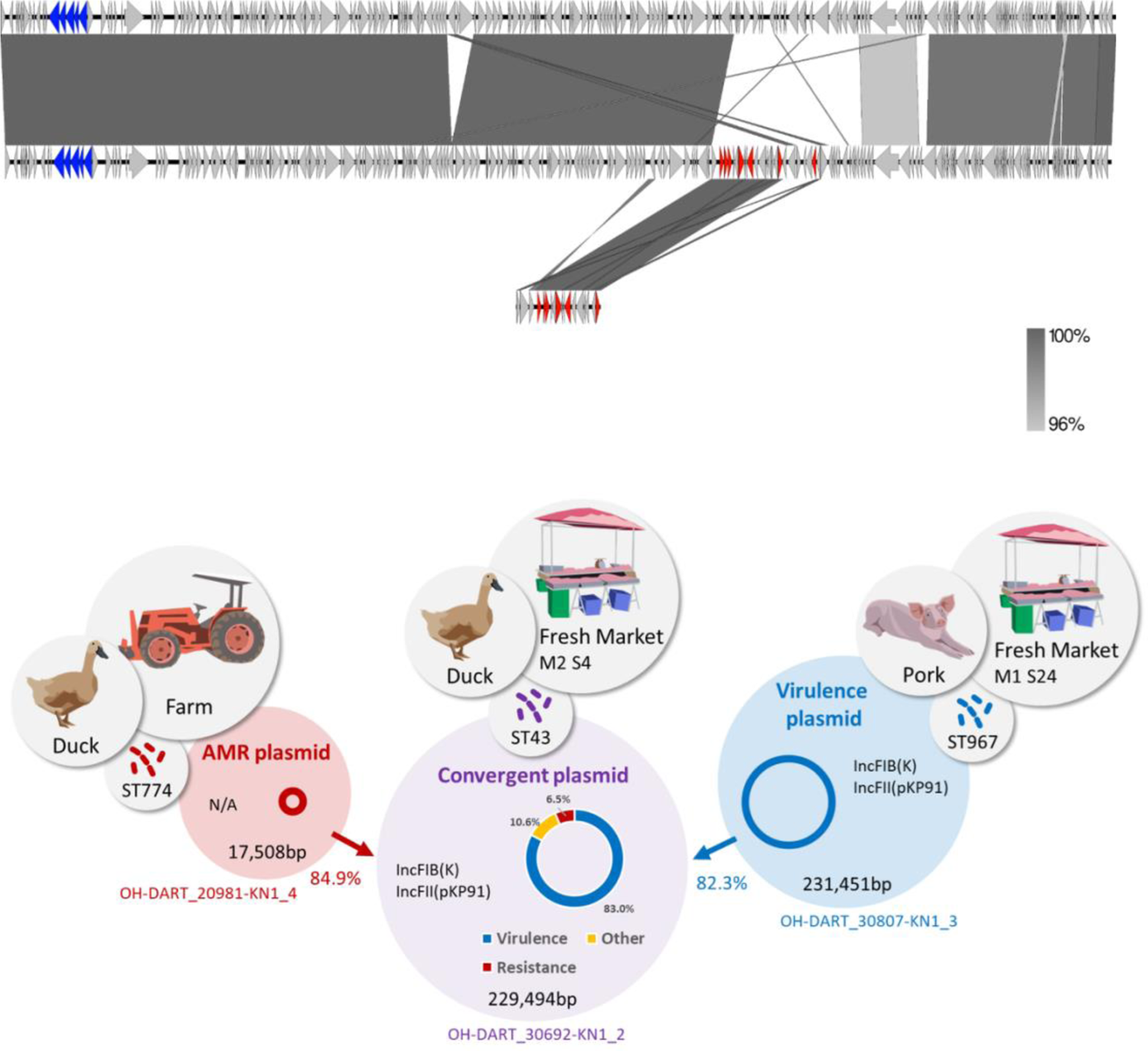
The convergent plasmid OH-DART_30692-KN1_2 was carried by an ST43 isolate (middle). This plasmid was recovered from duck meat and is highly similar to the *iuc*3 plasmid OH-DART_30807-KN1_3 isolated from pork meat (top). 83% of plasmid OH-DART_30692-KN1_2 shares >98% sequence identity with 82% of plasmid OH-DART_30807-KN1_3. The convergent plasmid OH-DART_30692-KN1_2 has incorporated 85% (>98% sequence identity) of the small (17508-bp) non-mobilizable plasmid OH-DART_20981-KN1_4 (bottom). This plasmid does not contain any rep genes and has the ARG profile *strA*.v1^; *strB*.v1; *floR*.v1; *sul2*; *tet*(A).v2; *bla*_SHV-12_. The convergent plasmid shares the same ARG profile, except with an additional *mph*A gene. The example provides a link between duck meat on the market, and samples from the neighbouring duck farm.

**Supplementary Figure 5C (Event 6).**
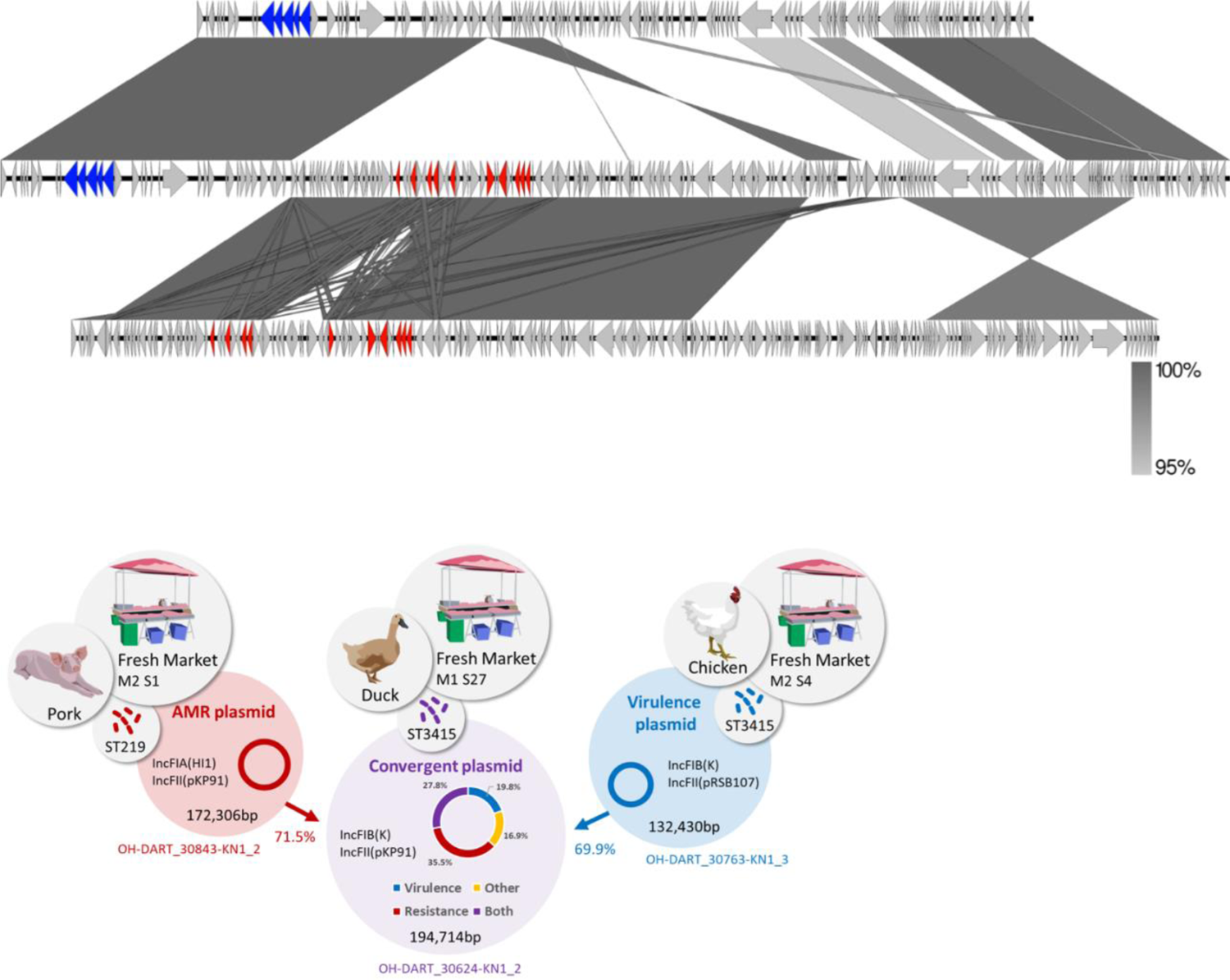
The convergent plasmid (OH-DART_30624-KN1_2) (middle) and both putative parental plasmids originate from the markets. 47.5% of the convergent plasmid shows a high level of nucleotide identity (>98%) to 69.9% putative *iuc*3 plasmid parent OH-DART_30763-KN1_3 (top), and these two plasmids were found in the same clone (ST3415). The putative AMR parental plasmid OH-DART_30843-KN1_2 (ST219) (bottom) shares an identical ARG profile with the convergent plasmid: *aac(3)-IId*^; *aadA17*\*; *strA*.v1^; *strB*.v1; *qnrS1*; *lnuF*.v1; *floR*.v2*; *sul2*; *tet*(A).v2; *bla*_SHV-12_. 71.5% of this plasmid shows a high nucleotide identity (98%) with 47.5% of the convergent plasmid. This plasmid was also identified as the putative AMR parental plasmid in Event 2. This example is noteworthy as a high percentage of the convergent plasmid (27.8%) shows a high level of nucleotide identity to both putative parents, reflecting the fact that the parents themselves share similar blocks of sequences.

**Supplementary Figure 5D (Event 7).**
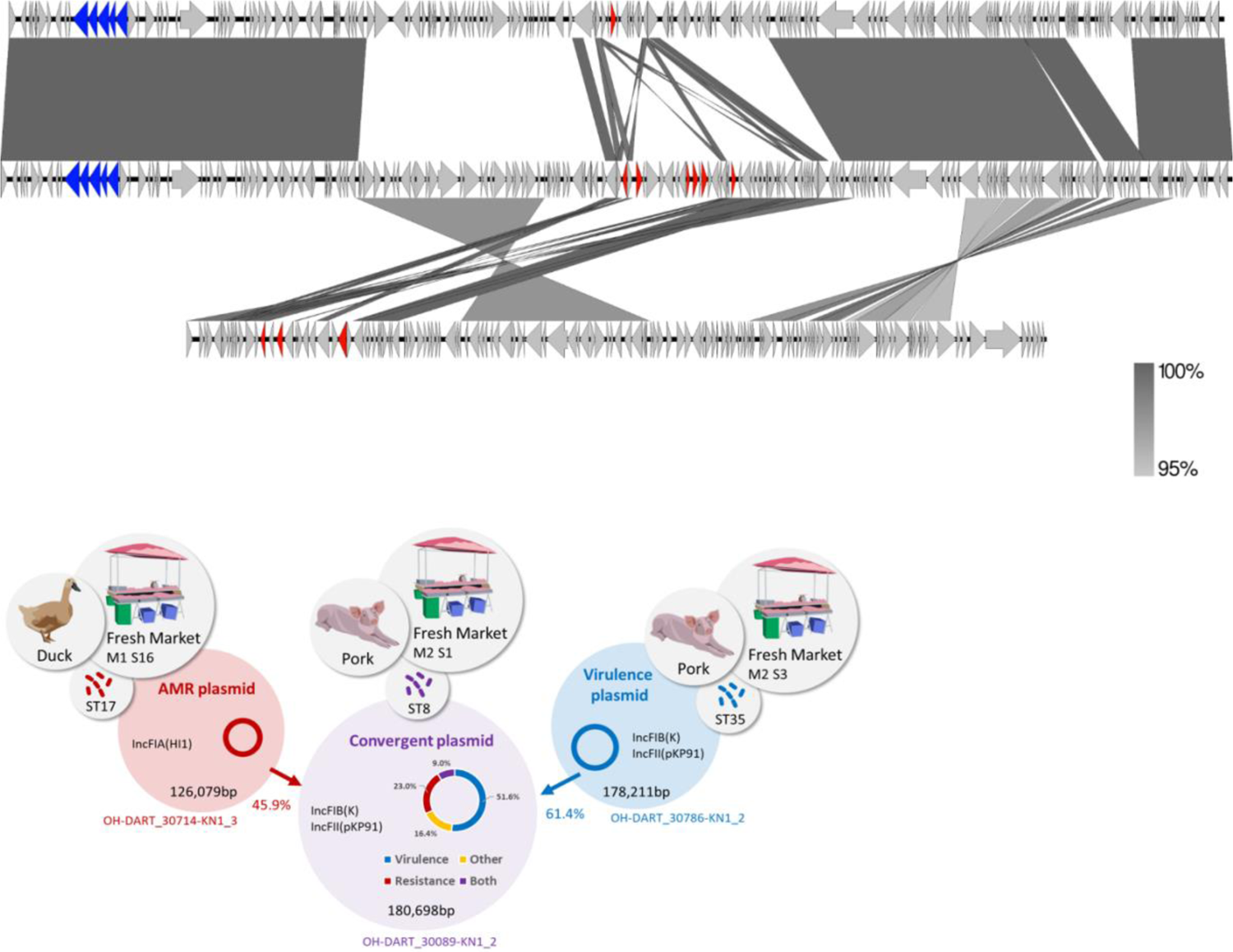
The convergence plasmid OH-DART_30089-KN1_2 (ST78) (middle) and both putative parents were isolated from the markets. There is a typically strong match between the convergence plasmid and the putative *iu*c3 parent OH-DART_30786-KN1_2; 60.6% of the convergence plasmid showed >98% nucleotide identity with 61.4% of this plasmid. However, in his case we were not able to identify a closely matching AMR parental plasmid. The ARG profile of the convergence plasmid is *aadA2*^ *qnrS1*; *catII*.2*; *sul1*; *dfrA12*; *bla*_CTX-M-55_. There are only 24 examples of Kp isolates harbouring *bla*_CTX-M-55_ in the OH-DART dataset (32 including other *Klebsiella* species). However, *bla*_CTX-M-55_ is very common among *E. coli* isolates. The closest match we could find among the fully closed assemblies was plasmid OH-DART_30714-KN1_3 (ST17); 45.9% of this plasmid shares high nucleotide identity (>95%) with 32% of the convergent plasmid; however, the ARG profile is different and in particular this plasmid carries *bla*_CTX-M-63_ rather the *bla*_CTX-M-55_. The ARG profile of OH-DART_30714-KN1_3 is: *qnrS1*; *tet*(A).v1^; *bla*_LAP-2_; *bla*_CTX-M-63_.

**Supplementary Figure 6.**
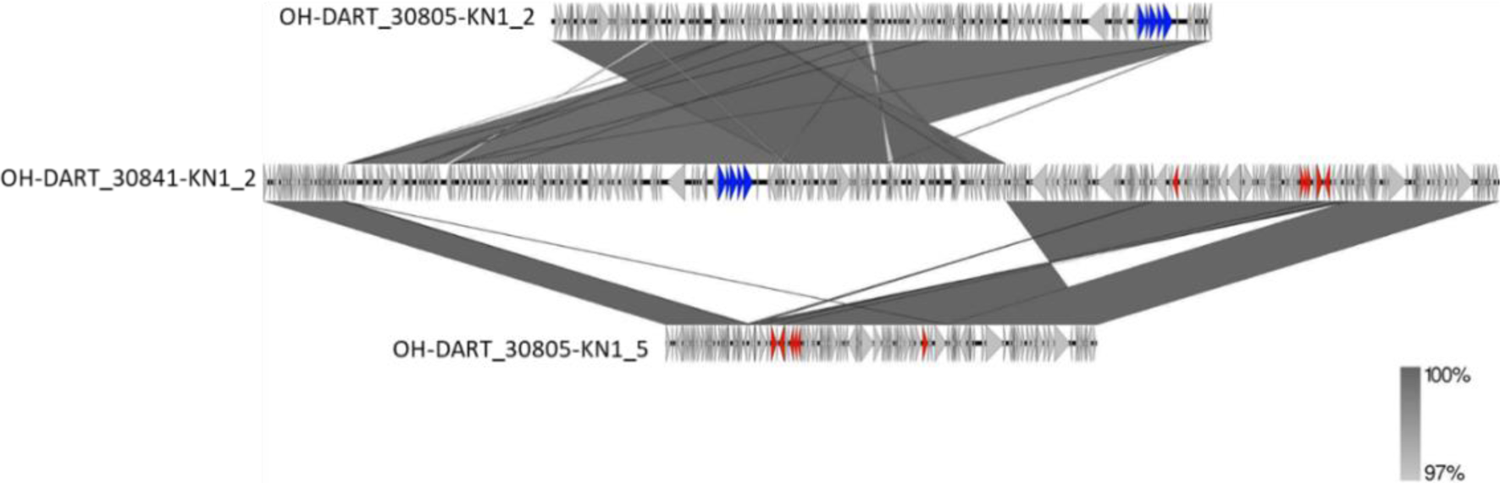
Alignment of three plasmids from Kp ST7513-1LV. The *iuc3* locus is in blue, and ARGs are in red. The top (*iuc*3) and bottom (AMR) plasmids were harboured by the same ST7513-1LV isolate (OH-DART_30805-KN1). Plasmid OH-DART_30805-KN1_2 has four replicon types: ColRNAI, IncFIB(AP001918), IncFIB(K)(pCAV1099-114), IncFII(pKP91), whilst plasmid OH-DART_30805-KN1_5 has only one (IncC). These have hybridised to form the middle large plasmid that was harboured by a different ST7513-1LV isolate (OH-DART_30841-KN1_2) and which contains all 5 replicon types (ColRNAI, IncC, IncFIB(AP001918), IncFIB(K)(pCAV1099-114), IncFII(pKP91)) and an identical ARG profile to OH-DART_30805-KN1_5 (*strAB*, *floR.v2\**, *sul2*, *tet*(A).v2, *bla*_CMY-2.v2_). Both isolates were sampled from the neighbouring Thai markets.

**Supplementary Figure 7.**
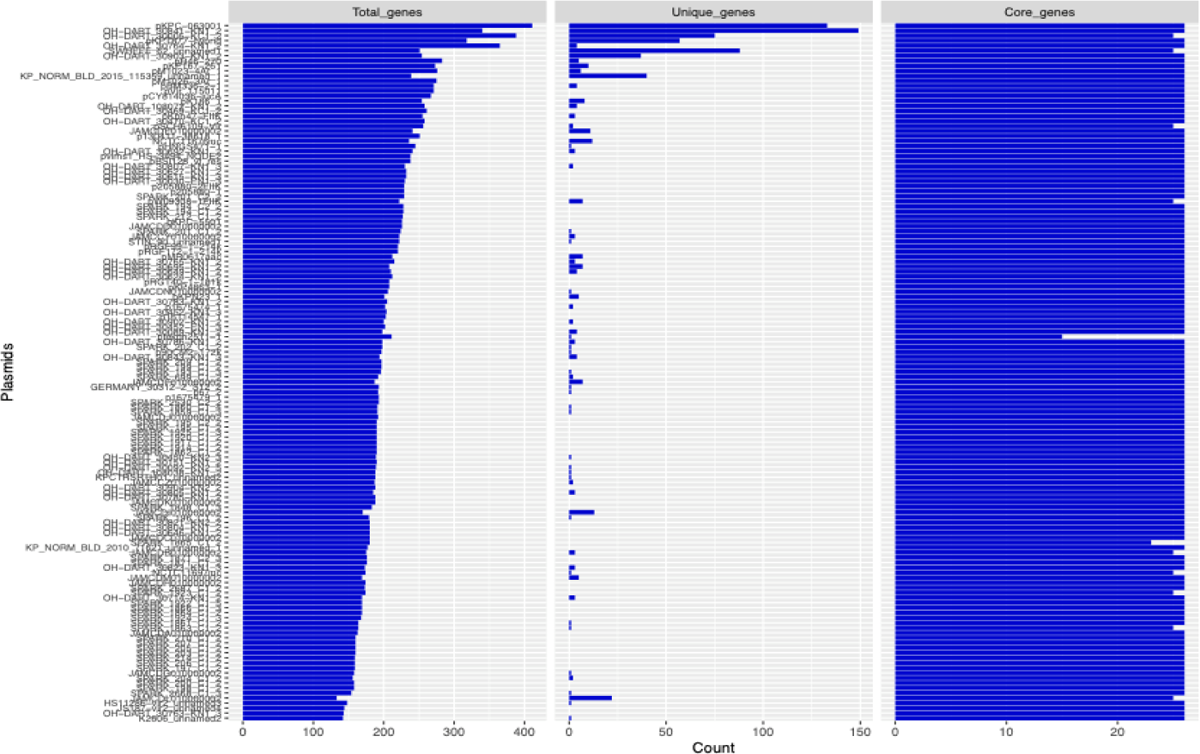
Number of total, unique and core genes identified by Roary in each of the 139 *iuc3*-carrying plasmids for which fully closed sequences are available.

**Supplementary Figure 8.**
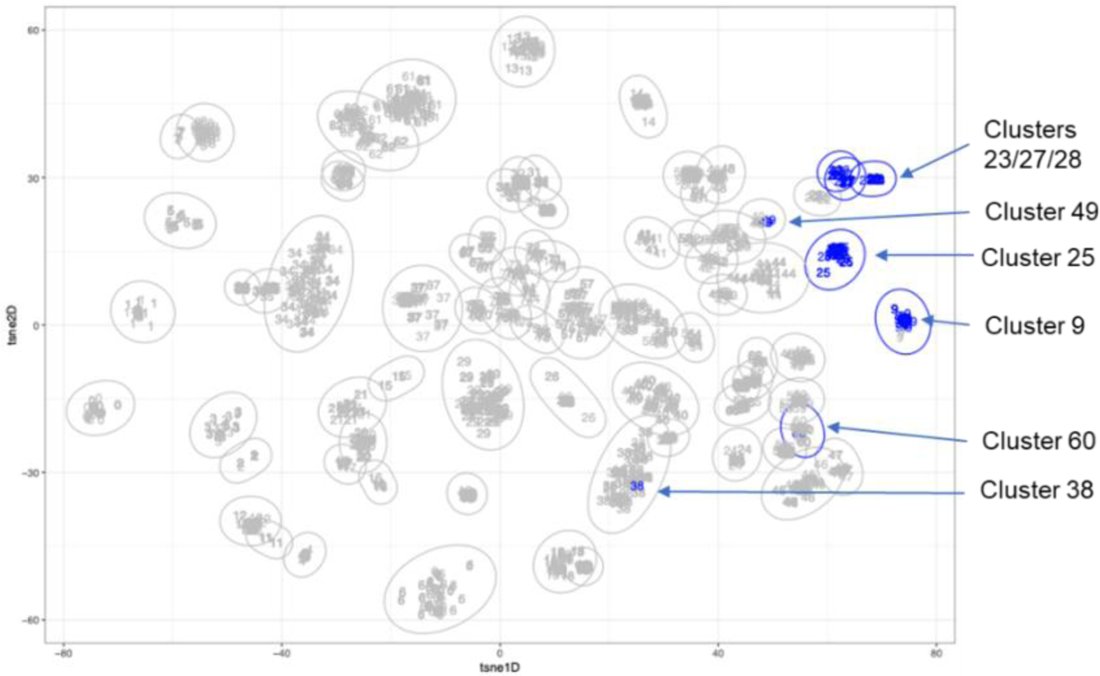
Clustering of 139 plasmids carrying *iuc*3 with 2874 plasmids from Thailand and Italy based on the mge-cluster analyses. Blue numbers and clusters represent plasmids carrying *iuc3*.

**Supplementary Figure 9.**
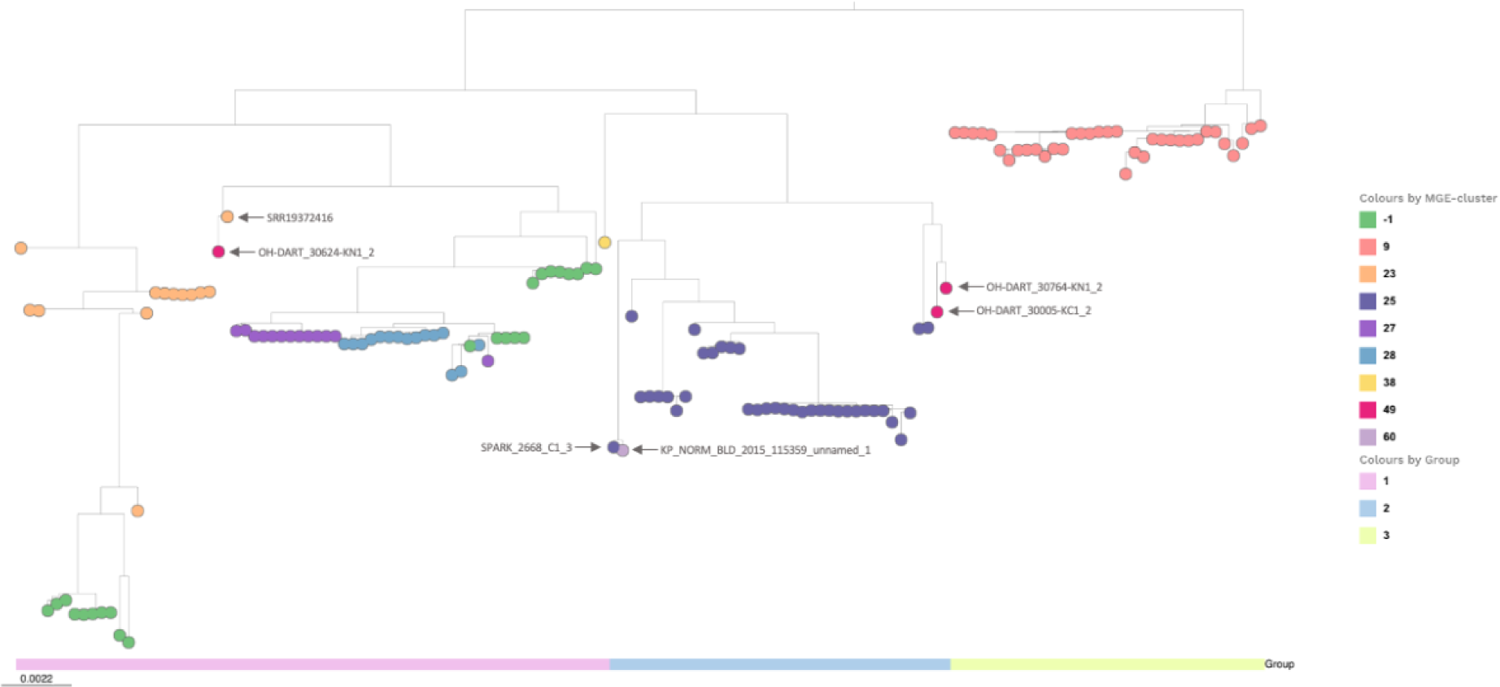
SNP tree of 139 *iuc*3 plasmid sequences. The colours of the nodes correspond to the cluster assignment by mge-cluster (Supplementary Figure 8). Clusters #9 and #25 are strongly supported by the tree and are assigned as Group 3 and Group 2 respectively. Group 1 represents a mixture of clusters. The clusters defined by mge-cluster are broadly consistent with a SNP-based tree constructed using complete plasmid genomes, with the following exceptions. Three convergent plasmids correspond to cluster #49, whilst on the tree OH-DART_30624-KN1_2 is placed in group 1, OH-DART_30005-KC1_2 and OH-DART_30764-KN1_2 are placed in group 2. OH-DART_30624-KN1_2 is positioned closely on the tree to the Norwegian porcine plasmid SRR19372416, which corresponds to cluster #23. KP_NORM_BLD_2015_115359_unnamed_1 (cluster #60) is positioned very closely on the tree to SPARK_2668_C1_3 (cluster #25). Of the nine convergent plasmids in total (including two from the public data), four correspond to Group 3 (cluster #9), whilst the other five are scattered across the tree.

**Supplementary Figure 10.**
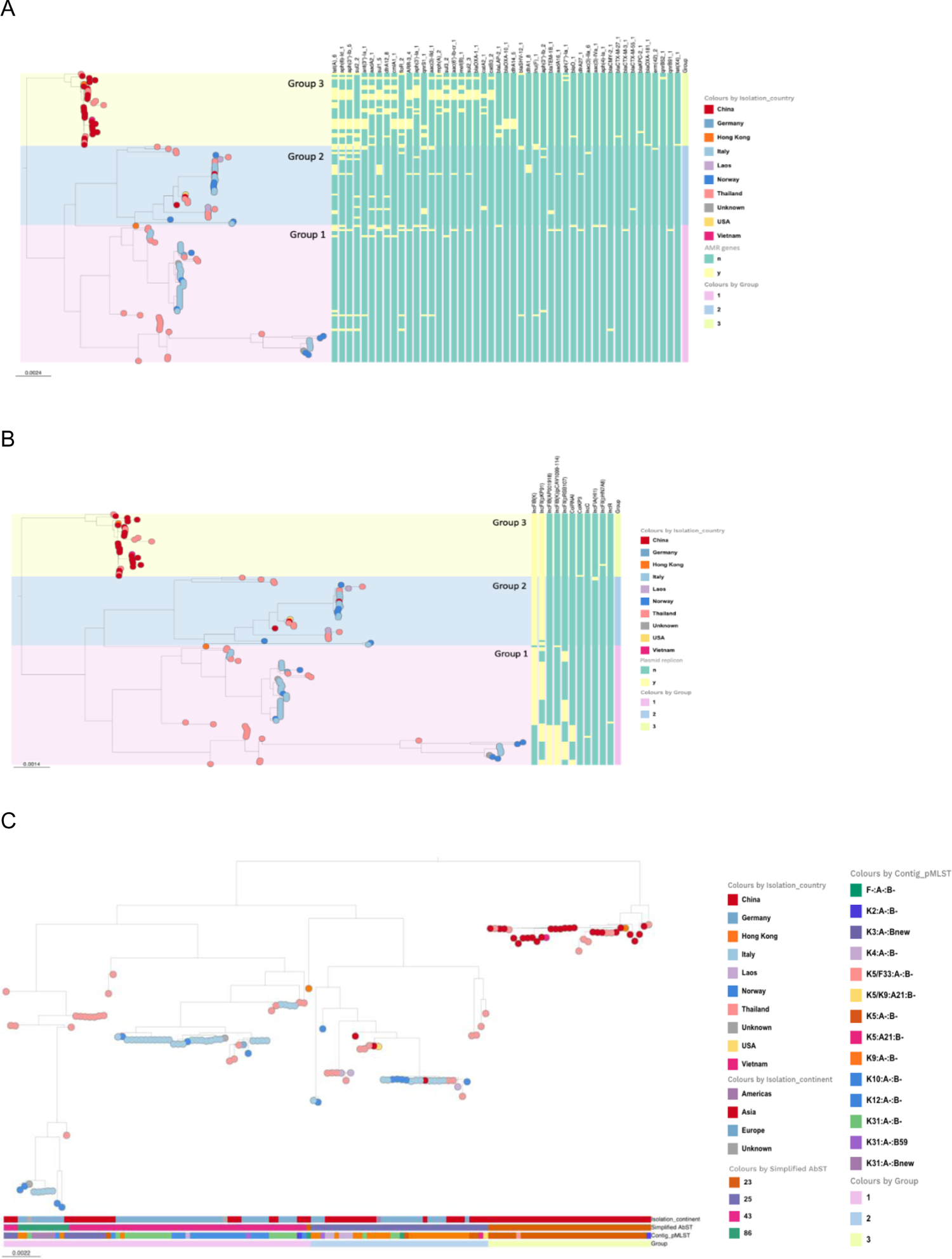
SNP tree of 139 *iuc*3 plasmids showing (**A**) ARGs (Resfinder), (**B**) replicon types and (**C**) geography, pMLST and simplified AbST.

**Supplementary Figure 11.**
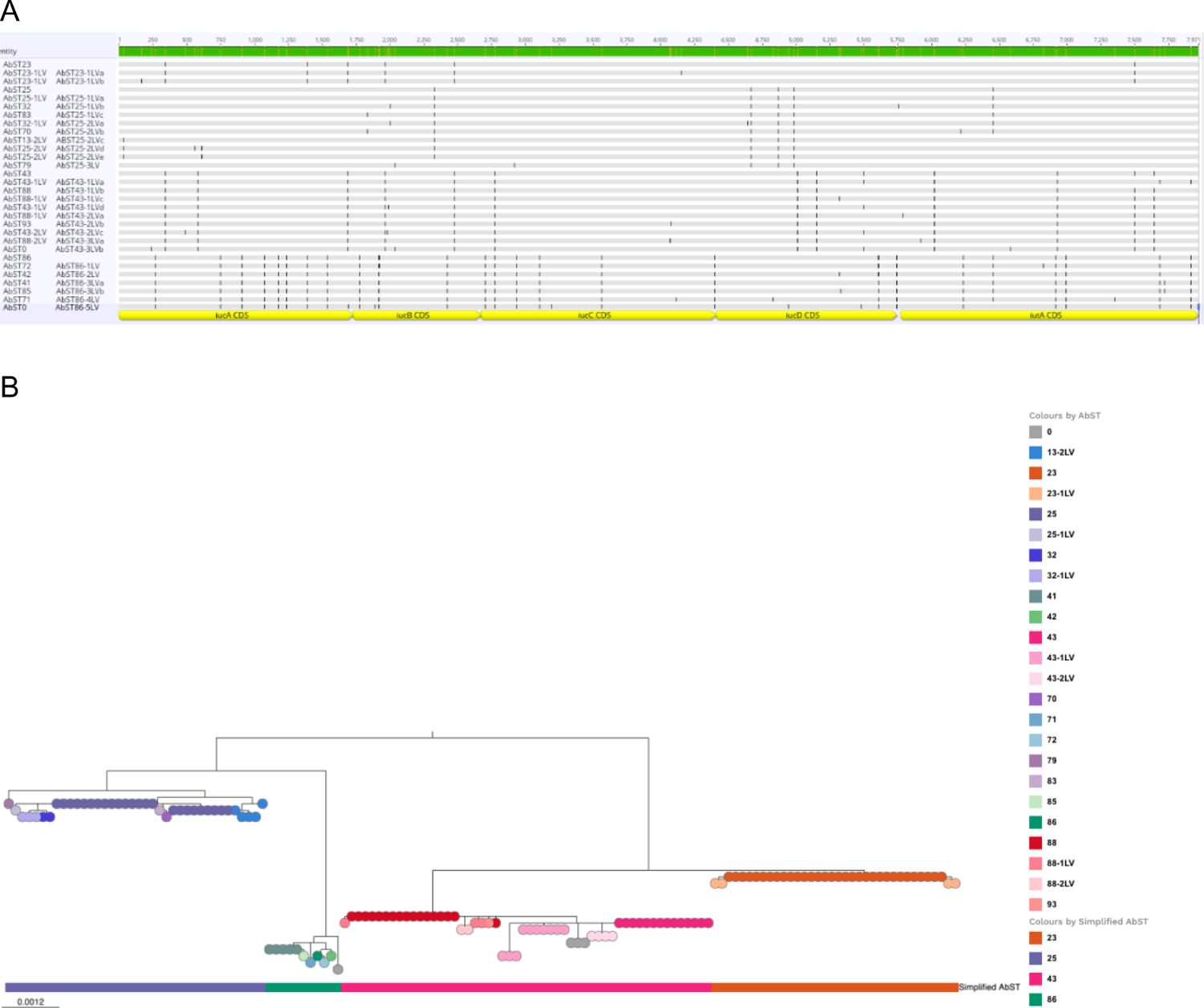
A. Alignment of the *iuc*3 locus in the 30 unique AbST types. Sequences are labelled on the left with the AbST assigned by Kleborate followed by the AbST assigned in this study (where different). The genes in the *iuc* locus are shown below the alignment. **B.** Phylogenetic tree based on the variation in the *iuc*3 locus within 139 complete *iuc*3 plasmid sequences. The colours on the nodes indicate the AbST assignments by Kleborate. These resolve into four distinct groups, assigned the name of the predominant AbST within the group, AbSTs 23, 25, 43 and 86, as indicated by the bar.

**Supplementary Figure 12.**
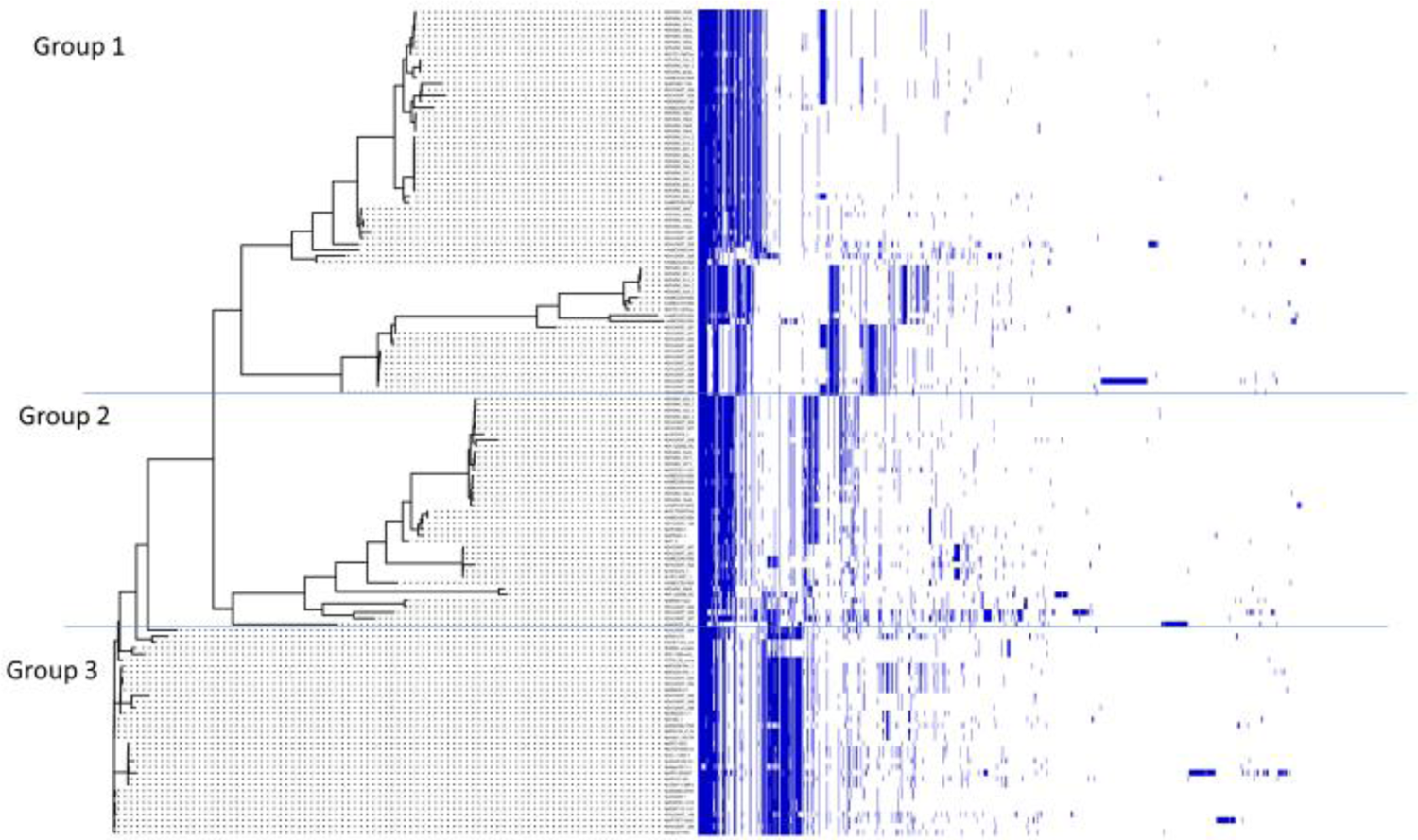
Gene content differences between the 139 *iuc*3 plasmids broken down by group. Genes are shown as blue bars in order of frequency. The tree on the left is based on SNPs.

**Supplementary Figure 13.**
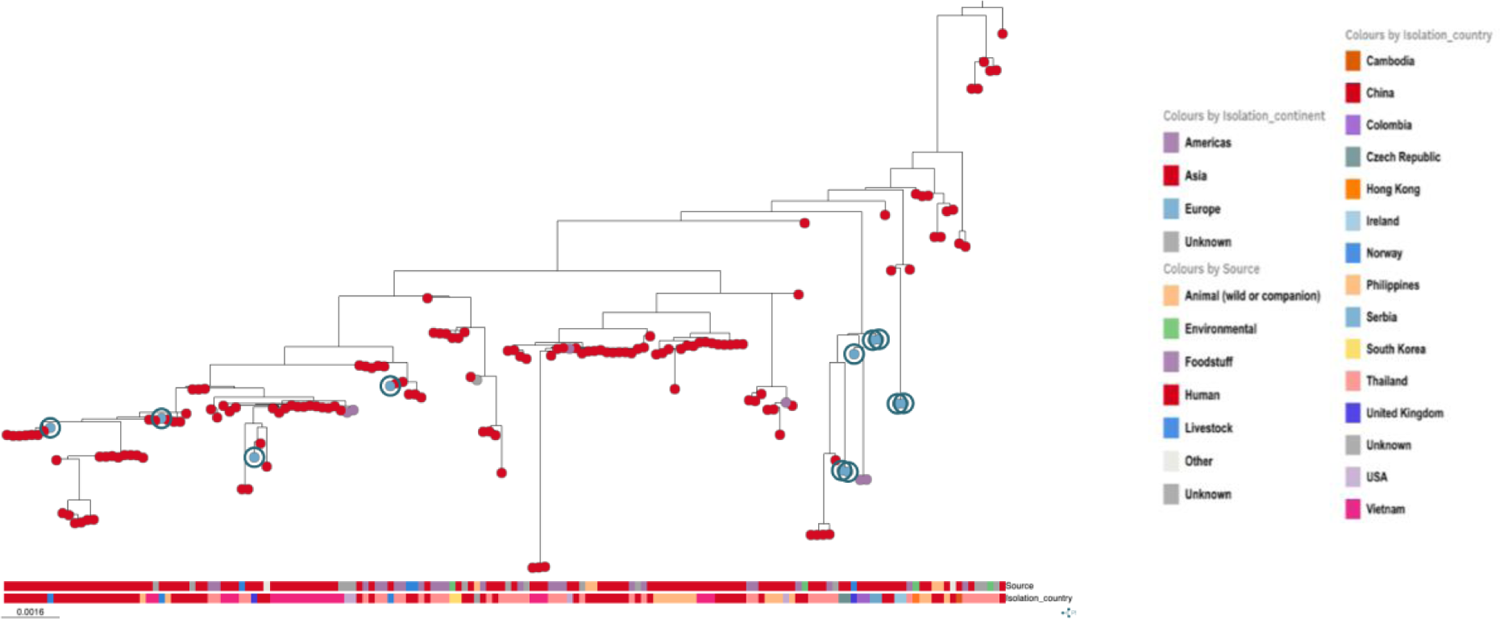
Group 3 subtree of the SNP-based tree of 517 plasmids. The nodes are coloured according to continent of origin, with European plasmids (blue) highlighted with a circle. The top bar gives the source and the bottom bar gives the country of origin. Ten of the eleven European plasmids were associated with humans, at least eight of them were from clinical isolates.

**Supplementary Figure 14.**
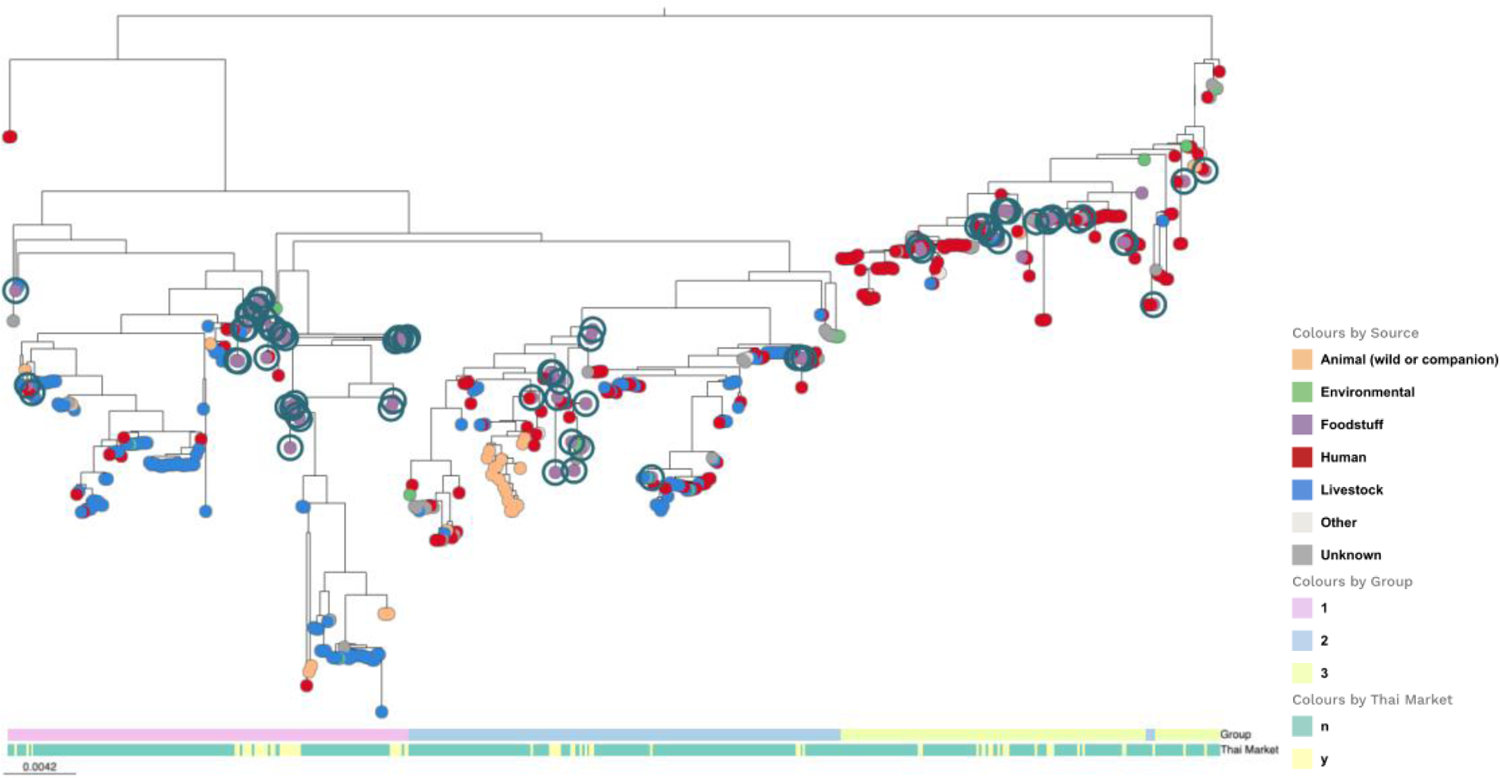
Tree of 517 *iuc*3 plasmids coloured by source. Plasmids derived from the two Thai markets (‘Foodstuffs’) are indicated by circles at the bottom bar and are scattered across the tree. The top bar indicates the plasmid group.

**Supplementary Figure 15.**
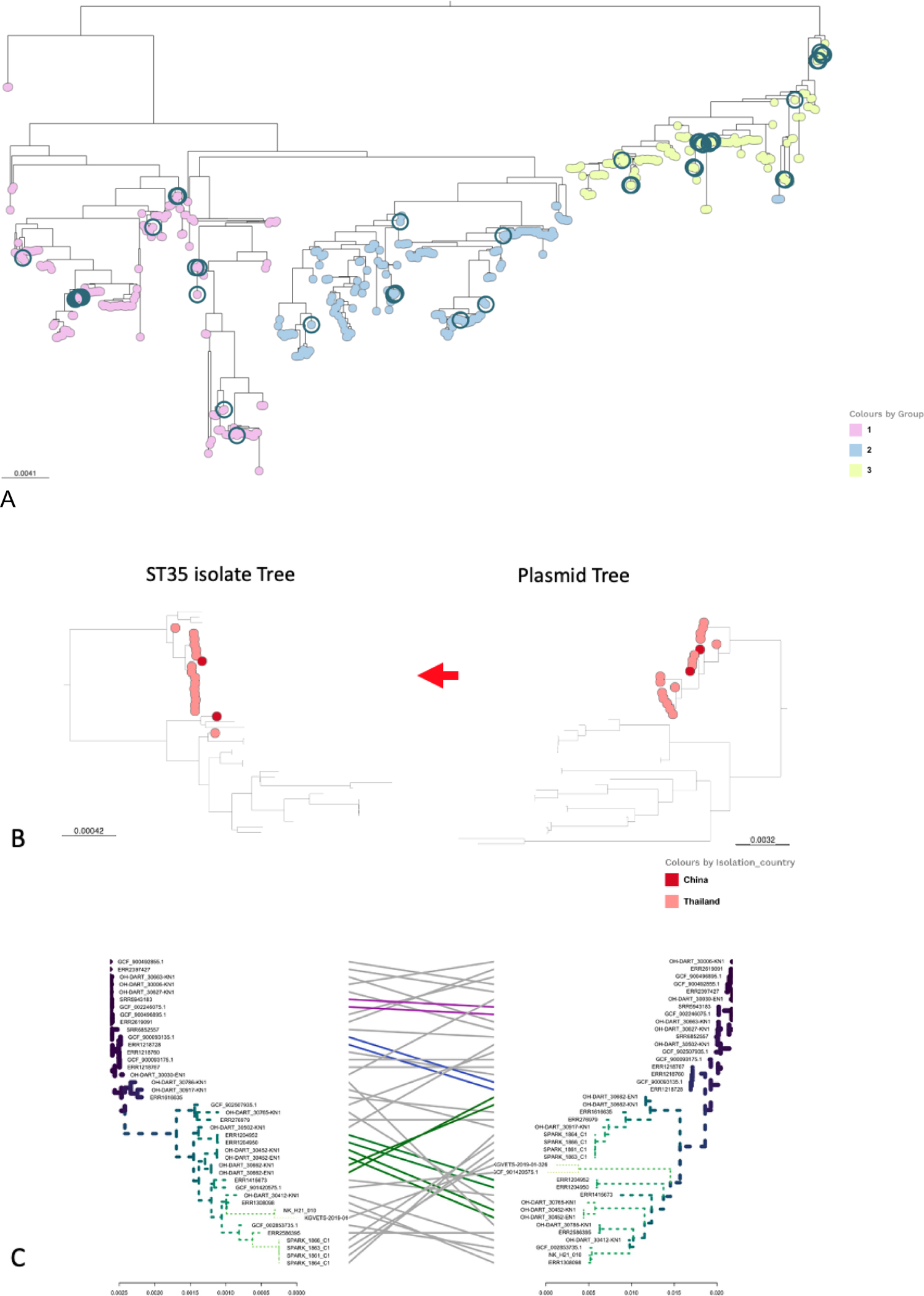
Analysis of ST35 isolates and plasmids **A.** The tree of 517 *iuc*3 plasmids with those harboured by ST35 isolates highlighted. The node colours indicate Groups. **B.** A comparison of the tree of all 41 ST35 isolates harbouring an *iuc*3 plasmid, and the tree of the plasmids in these isolates. The group 3 plasmids are highlighted on the plasmid tree, the colour of the node indicates country of origin. The corresponding isolates are highlighted on the isolates tree. The group 3 plasmids are found in a distinct sub lineage of ST35, with two exceptions. There is little consistency between the two trees when the other plasmids are considered. **C.** A tanglegram of the two trees, lines connect isolates and their corresponding plasmids.

## Supplementary Tables

**Supplementary Table 1.**
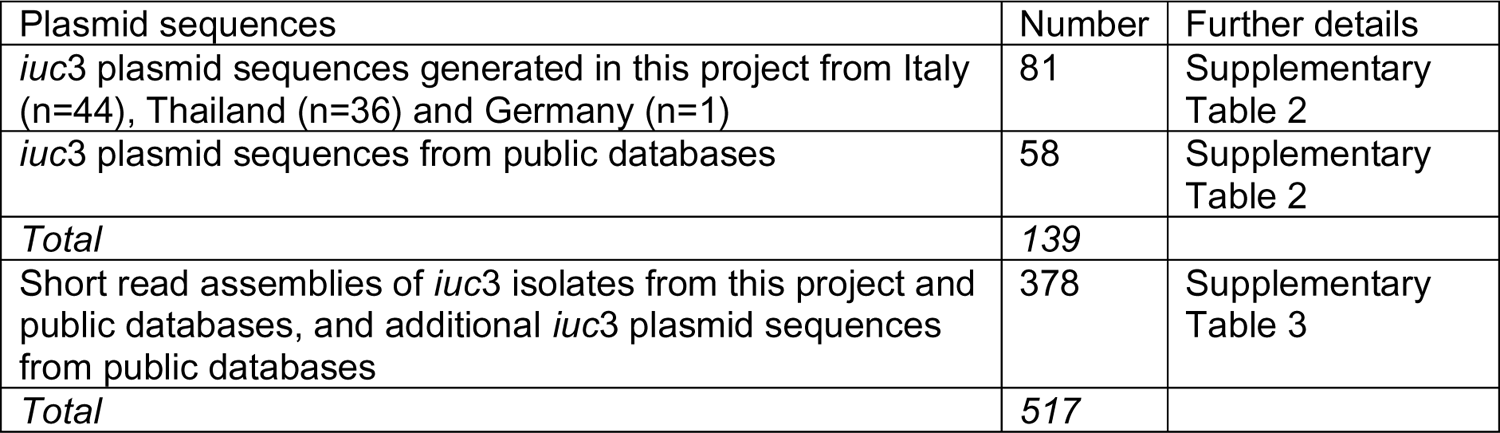
Data summary of 139 long-read *iuc*3 plasmid sequences, 81 generated in this project and 58 from public databases which were used for initial analysis, and subsequently combined with 378 further short-read sequences from *iuc*3 isolates.

| Plasmid sequences | Number | Further details |
| --- | --- | --- |
| *iuc3* plasmid sequences generated in this project from Italy (n=44), Thailand (n=36) and Germany (n=1) | 81 | Supplementary Table 2 |
| *iuc3* plasmid sequences from public databases | 58 | Supplementary Table 2 |
| *Total* | 139 |  |
| Short read assemblies of *iuc3* isolates from this project and public databases, and additional *iuc3* plasmid sequences from public databases | 378 | Supplementary Table 3 |
| *Total* | 517 |  |

**Supplementary Tables 2, 3** and 4 are provided as separate files.

**Supplementary Table 4.** Frequency of IS families in 139 *iuc*3 plasmids based on annotations provided by Prokka.

|  | No. of sequences |
| --- | --- |
| IS3 family | 562 |
| IS110 family | 297 |
| IS481 family | 257 |
| IS6 family | 180 |
| ISNCY family | 131 |
| IS1 family | 110 |
| IS200/IS605 family | 97 |
| ISL3 family | 78 |
| IS5 family | 73 |
| IS21 family | 54 |
| IS66 family | 36 |
| IS256 family | 30 |
| ISKra4 family | 16 |
| IS1595 family | 15 |
| IS150 family | 12 |
| IS1380 family | 6 |
| IS91 family | 6 |
| IS4 family | 4 |
| IS630 family | 3 |
| IS1182 family | 2 |
| IS30 family | 1 |

**Supplementary Table 5.** SNP distances between the four simplified AbST groupings of *iuc*3 in the 139 *iuc*3-positive assemblies (Supplementary Table 1). Multiple AbST variants have been simplified into each grouping (Supplementary Methods and Supplementary Figure 11).

|  | AbST 23 | AbST 25 | AbST 43 | AbST 86 |
| --- | --- | --- | --- | --- |
| AbST 23 | 0 - 2 | 11 - 14 | 8 - 13 | 33 - 37 |
| AbST 25 | 11 - 14 | 0 - 6 | 17 - 23 | 30 - 37 |
| AbST 43 | 8 - 13 | 17 - 23 | 0 - 8 | 35 - 46 |
| AbST 86 | 33 - 37 | 30 - 37 | 35 - 46 | 0 - 9 |

**Supplementary Table 6.**
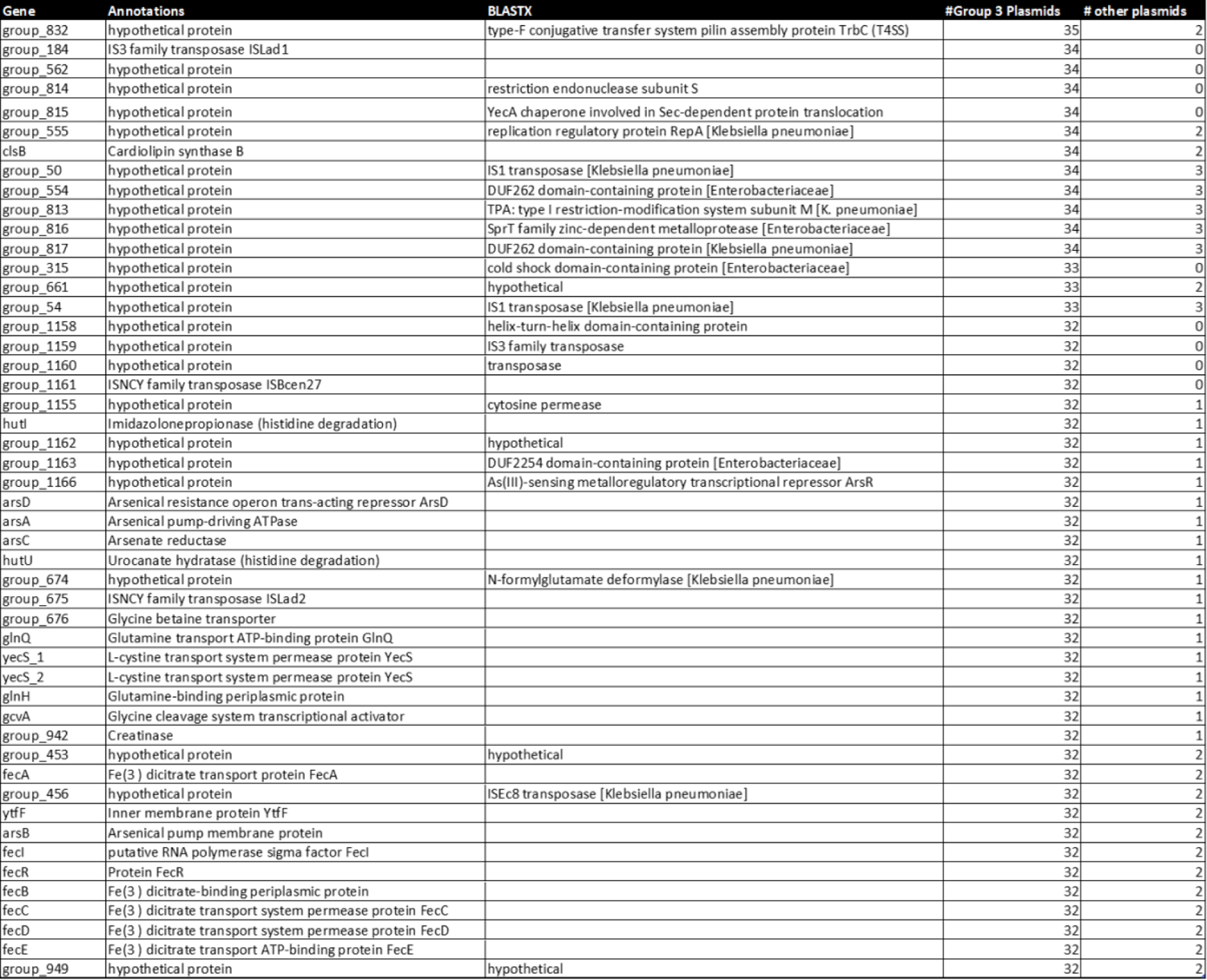
Genes associated with the 139 *iuc*3 plasmids assigned to Group 3 (n=35). The Prokka annotation / Roary assignment is shown in the first column. All genes assigned as hypothetical by Prokka were checked using BLASTX and the best hits are given in the 3rd column.

**Supplementary Table 7.** The geographical and ecological distribution of the 517 plasmids according to Group. The closely related Thai plasmids pMR0617aac, GCF_002752955.1 and 480738_p1 (from faecal flora from hospitalised patients) and OH-DART_30092-KN2 (from pork meat) were assigned as Group 1 based on the tree of 139 plasmids and the AbST data, rather than Group 3 as indicated by the tree of 517 plasmids (see text).

|  |  | Group 1 | Group 2 | Group 3 |
| --- | --- | --- | --- | --- |
| Source | Animal | 111 (64.9 %) | 76 (40.4 %) | 8 (5 %) |
|  | Environment | 4 (2.3 %) | 8 (4.3 %) | 4 (2.5 %) |
|  | Foodstuff | 33 (19.3 %) | 17 (9.0 %) | 23 (14.6 %) |
|  | Human | 16 (9.4 %) | 66 (35 %) | 104 (65.8 %) |
|  | Other / Unknown | 7 (4.1 %) | 21 (11.2 %) | 19 (12.0 %) |
| Continent | Africa | 4 (2.3 %) | 3 (1.6 %) | 0 |
|  | Americas | 2 (1.2 %) | 8 (4.3 %) | 6 (3.8 %) |
|  | Asia | 45 (26.3 %) | 81 (43.1 %) | 140 (88.6 %) |
|  | Australasia | 3 (1.8 %) | 0 | 0 |
|  | Europe | 113 (66.1 %) | 96 (51.1 %) | 11 (7.0 %) |
|  | Unknown | 4 (2.3 %) | 0 | 1 (0.6 %) |
|  | Total | 171 | 188 | 158 |

## Supplementary Methods

### OH-DART sampling

All sampling was carried out in a defined semi-urban region, Bang Len, in Nakhon Pathom province, central Thailand. Over 7,000 samples from hospital patients, community carriage, agriculture, farms, food (markets), soil and water were taken from October 2019 to March 2021. Samples were initially screened using Chromagar 3GC-R or Chromagar CPE, and selected colonies were re-streaked on Brilliance Selective Medium (ThermoFisher Scientific) with either cefotaxime 2 µg/ml or ertapenem 0.5 µg/ml. A total of 607 Kp isolates were recovered from selective media (either with third generation cephalosporin or carbapenem) and selected for Illumina sequencing. 591 high quality Kp genomes were generated. It was not possible to sample from pig farms due to an outbreak of African Swine Fever in the region.

### Sequencing of isolates from Thailand

A single colony of each isolate was picked from a fresh SCAI plate containing ampicllin (10 µg/ml) into LB broth (Miller) with ampicillin (10 μg/ml) and incubated at 37°C overnight with shaking. For short read sequencing, DNA was extracted using a Monarch genomic DNA purification kit (New England Biolabs) and quantified with the Qubit 4.0 system (Thermo Fisher) or with an in-house method at MicrobesNG (https://microbesng.com/). Isolates were sequenced using Illumina sequencers (HiSeq/NovaSeq) using a 250-bp paired-end protocol. Short-reads were trimmed using Trimmomatic v0.30^1^ and the trimmed reads were used to generate *de novo* assemblies using SPAdes v3.7 ^2^. Long read sequences were generated from DNA extracted with an in-house method at MicrobesNG, using a GridION (Oxford Nanopore Technologies, Oxford, UK).

### Bioinformatic analysis of plasmids

Mash distances and trees were generated using mashtree v1.2.0 ^3^. To generate a SNP-based plasmid tree, we first used Snippy v4.6.0 (https://github.com/tseemann/snippy) using with the assembled plasmid files as input, and the largest closed *iuc*3 plasmid in the data (OH-DART_30005-KC1_2) as reference.

Phylogenetic analysis was carried out using FastTree v.2.1.11 on the whole genome SNP alignment file ^4,5^. Roary was used to identify the core and accessory genes of the 139 plasmids ^6^. The plasmids were analysed using Kleborate v2.3.2, ABRicate v.1.0.1 with the resfinder (version 2023-Oct-31) and plasmidfinder (version 2021-Mar-27) databases and pMLST (https://bitbucket.org/genomicepidemiology/pmlst/src/master/). Mob-typer (from MOB-suite v3.1.5) was used to predict plasmid mobility ^7^. The trees were combined with metadata and output from Kleborate, ABRicate and pMLST and visualised using Microreact v233^8^. Closed plasmids from the entire SpARK and OH-DART collections were analysed as implemented in MGE-cluster using default parameters^9^.

### Simplified AbST scheme

The aerobactin locus has five genes, each of which can occur as one of several alleles. Some of the combinations of alleles have been designated AbSTs, and variants of these are referred to as AbSTX-1LV, etc^10^.

We used Kleborate to call the alleles and assign AbSTs to the 139 *iuc*3-positive plasmid assemblies (Supplementary Table 2), revealing 30 unique AbST types. An alignment and SNP-based phylogenetic tree of the locus extracted from the assemblies (Supplementary Figure 11) revealed that they cluster into four groups, which we have named for the predominant AbST within the group. We refer to these as simplified AbSTs.

## Supplementary Note

### AbST25 plasmids in group 3

In the SNP tree of 517 *iuc*3 isolates (Figure 3A), 4 plasmids with simplified AbST25 appear on a branch within Group 3, whereby all other plasmids in this group correspond to simplified AbST23. Two of these plasmids (pMR0617aac and OH-DART_30092-KN2) are also present in the tree of 139 plasmids, where they are positioned within Group 2 (Supplementary Figure 10c). The most likely explanation therefore is that the position of these four plasmids on the tree in Figure 3 is artefactual. These plasmids are closely related and positioned on the end of a relatively long branch. Three of them were harboured by clinical ST45 isolates from Thailand (pMR0617aac;^11^ GCF_002752955.1 and 480738_p1), with the third from an ST520 isolate from pork meat from the OH-DART study (OH-DART_30092-KN2). The similarity of these three plasmids thus acts as further evidence for plasmid movement between foodborne and clinical isolates in Thailand. The SWHEFF_62 plasmid, which clusters atypically by MGE-cluster, corresponds to AbST23 but does not cluster with group 3. This is consistent with the transfer of the *iuc*3 locus into a distinct plasmid group.

## References

1 Tacconelli E, Carrara E, Savoldi A, et al. Discovery, research, and development of new antibiotics: the WHO priority list of antibiotic-resistant bacteria and tuberculosis. Lancet Infect Dis 2018; 18: 318–27.

2 Rello J, Kalwaje Eshwara V, Lagunes L, et al. A global priority list of the TOp TEn resistant Microorganisms (TOTEM) study at intensive care: a prioritization exercise based on multi-criteria decision analysis. Eur J Clin Microbiol Infect Dis 2019; 38: 319– 23.

3 Russo TA, Marr CM. Hypervirulent Klebsiella pneumoniae. Clin Microbiol Rev 2019; 32. DOI:10.1128/CMR.00001-19.

4 Wyres KL, Lam MMC, Holt KE. Population genomics of Klebsiella pneumoniae. Nat Rev Microbiol 2020; 18: 344–59.

5 Wyres KL, Nguyen TNT, Lam MMC, et al. Genomic surveillance for hypervirulence and multi-drug resistance in invasive Klebsiella pneumoniae from South and Southeast Asia. Genome Med 2020; 12: 11.

6 David S, Cohen V, Reuter S, et al. Integrated chromosomal and plasmid sequence analyses reveal diverse modes of carbapenemase gene spread among Klebsiella pneumoniae. Proc Natl Acad Sci U S A 2020; 117: 25043–54.

7 Marr CM, Russo TA. Hypervirulent Klebsiella pneumoniae: a new public health threat. Expert Rev Anti Infect Ther 2019; 17: 71–3.

8 Chen L, Kreiswirth BN. Convergence of carbapenem-resistance and hypervirulence in Klebsiella pneumoniae. Lancet Infect. Dis. 2018; 18: 2–3.

9 Xia P, Yi M, Yuan Y, et al. Coexistence of Multidrug Resistance and Virulence in a Single Conjugative Plasmid from a Hypervirulent Klebsiella pneumoniae Isolate of Sequence Type 25. mSphere 2022; 7: e0047722.

10 Lam MMC, Wick RR, Watts SC, Cerdeira LT, Wyres KL, Holt KE. A genomic surveillance framework and genotyping tool for Klebsiella pneumoniae and its related species complex. Nat Commun 2021; 12: 4188.

11 Yang X, Dong N, Chan EW-C, Zhang R, Chen S. Carbapenem Resistance-Encoding and Virulence-Encoding Conjugative Plasmids in Klebsiella pneumoniae. Trends Microbiol 2021; 29: 65–83.

12 Zhang Y, Jin L, Ouyang P, et al. Evolution of hypervirulence in carbapenem-resistant Klebsiella pneumoniae in China: a multicentre, molecular epidemiological analysis. J Antimicrob Chemother 2020; 75: 327–36.

13 Pei N, Li Y, Liu C, et al. Large-Scale Genomic Epidemiology of Klebsiella pneumoniae Identified Clone Divergence with Hypervirulent Plus Antimicrobial-Resistant Characteristics Causing Within-Ward Strain Transmissions. Microbiol Spectr 2022; 10: e0269821.

14 Gu D, Dong N, Zheng Z, et al. A fatal outbreak of ST11 carbapenem-resistant hypervirulent Klebsiella pneumoniae in a Chinese hospital: a molecular epidemiological study. Lancet Infect Dis 2018; 18: 37–46.

15 Zhou K, Xue C-X, Xu T, et al. A point mutation in recC associated with subclonal replacement of carbapenem-resistant Klebsiella pneumoniae ST11 in China. Nat Commun 2023; 14: 2464.

16 Li R, Cheng J, Dong H, et al. Emergence of a novel conjugative hybrid virulence multidrug-resistant plasmid in extensively drug-resistant Klebsiella pneumoniae ST15. Int J Antimicrob Agents 2020; 55: 105952.

17 Xie M, Dong N, Chen K, et al. A hybrid plasmid formed by recombination of a virulence plasmid and a resistance plasmid in Klebsiella pneumoniae. J Glob Antimicrob Resist 2020; 23: 466–70.

18 Lam MMC, Wyres KL, Wick RR, et al. Convergence of virulence and MDR in a single plasmid vector in MDR Klebsiella pneumoniae ST15. J Antimicrob Chemother 2019; 74: 1218–22.

19 Turton J, Davies F, Turton J, Perry C, Payne Z, Pike R. Hybrid Resistance and Virulence Plasmids in ‘High-Risk’ Clones of Klebsiella pneumoniae, Including Those Carrying blaNDM-5. Microorganisms 2019; 7. DOI:10.3390/microorganisms7090326.

20 Arcari G, Carattoli A. Global spread and evolutionary convergence of multidrug-resistant and hypervirulent Klebsiella pneumoniae high-risk clones. Pathog Glob Health 2023; 117: 328–41.

21 Shankar C, Vasudevan K, Jacob JJ, et al. Hybrid Plasmids Encoding Antimicrobial Resistance and Virulence Traits Among Hypervirulent Klebsiella pneumoniae ST2096 in India. Front Cell Infect Microbiol 2022; 12: 875116.

22 Ahmed MAE-GE-S, Yang Y, Yang Y, et al. Emergence of Hypervirulent Carbapenem-Resistant Klebsiella pneumoniae Coharboring a blaNDM-1-Carrying Virulent Plasmid and a blaKPC-2-Carrying Plasmid in an Egyptian Hospital. mSphere 2021; 6. DOI:10.1128/mSphere.00088-21.

23 Starkova P, Lazareva I, Avdeeva A, et al. Emergence of Hybrid Resistance and Virulence Plasmids Harboring New Delhi Metallo-β-Lactamase in Klebsiella pneumoniae in Russia. Antibiotics (Basel) 2021; 10. DOI:10.3390/antibiotics10060691.

24 Russo TA, Olson R, MacDonald U, Beanan J, Davidson BA. Aerobactin, but not yersiniabactin, salmochelin, or enterobactin, enables the growth/survival of hypervirulent (hypermucoviscous) Klebsiella pneumoniae ex vivo and in vivo. Infect Immun 2015; 83: 3325–33.

25 Lam MMC, Wyres KL, Judd LM, et al. Tracking key virulence loci encoding aerobactin and salmochelin siderophore synthesis in Klebsiella pneumoniae. Genome Med 2018; 10: 77.

26 Thorpe HA, Booton R, Kallonen T, et al. A large-scale genomic snapshot of Klebsiella spp. isolates in Northern Italy reveals limited transmission between clinical and non-clinical settings. Nat Microbiol 2022; 7: 2054–67.

27 Kaspersen H, Franklin-Alming FV, Hetland MAK, et al. Highly conserved composite transposon harbouring aerobactin iuc3 in Klebsiella pneumoniae from pigs. Microb Genom 2023; 9. DOI:10.1099/mgen.0.000960.

28 Klaper K, Hammerl JA, Rau J, Pfeifer Y, Werner G. Genome-Based Analysis of Klebsiella spp. Isolates from Animals and Food Products in Germany, 2013–2017. Pathogens 2021; 10: 573.

29 Booton RD, Meeyai A, Alhusein N, et al. One Health drivers of antibacterial resistance: Quantifying the relative impacts of human, animal and environmental use and transmission. One Health 2021; 12: 100220.

30 Van Kregten E, Westerdaal NA, Willers JM. New, simple medium for selective recovery of Klebsiella pneumoniae and Klebsiella oxytoca from human feces. J Clin Microbiol 1984; 20: 936–41.

31 Price MN, Dehal PS, Arkin AP. FastTree: computing large minimum evolution trees with profiles instead of a distance matrix. Mol Biol Evol 2009; 26: 1641–50.

32 Price MN, Dehal PS, Arkin AP. FastTree 2--approximately maximum-likelihood trees for large alignments. PLoS One 2010; 5: e9490.

33 Katz LS, Griswold T, Morrison SS, et al. Mashtree: a rapid comparison of whole genome sequence files. J Open Source Softw 2019; 4. DOI:10.21105/joss.01762.

34 Wick RR, Judd LM, Gorrie CL, Holt KE. Unicycler: Resolving bacterial genome assemblies from short and long sequencing reads. PLoS Comput Biol 2017; 13: e1005595.

35 Seemann T. Prokka: rapid prokaryotic genome annotation. Bioinformatics 2014; 30: 2068–9.

36 Argimón S, Abudahab K, Goater RJE, et al. Microreact: visualizing and sharing data for genomic epidemiology and phylogeography. Microb Genom 2016; 2: e000093.

37 Che Y, Xu X, Yang Y, et al. High-resolution genomic surveillance elucidates a multilayered hierarchical transfer of resistance between WWTP- and human/animal-associated bacteria. Microbiome 2022; 10: 16.

38 Yang QE, Tansawai U, Andrey DO, et al. Environmental dissemination of mcr-1 positive Enterobacteriaceae by Chrysomya spp. (common blowfly): An increasing public health risk. Environ Int 2019; 122: 281–90.

39 Wang X, Zhao J, Ji F, et al. Multiple-Replicon Resistance Plasmids of Klebsiella Mediate Extensive Dissemination of Antimicrobial Genes. Front Microbiol 2021; 12: 754931.

40 Wu F, Ying Y, Yin M, et al. Molecular Characterization of a Multidrug-Resistant Klebsiella pneumoniae Strain R46 Isolated from a Rabbit. Int J Genomics Proteomics 2019; 2019: 5459190.

41 Argimón S, David S, Underwood A, et al. Rapid Genomic Characterization and Global Surveillance of Klebsiella Using Pathogenwatch. Clin Infect Dis 2021; 73: S325–35.

42 Zeng L, Zhang J, Hu K, et al. Microbial Characteristics and Genomic Analysis of an ST11 Carbapenem-Resistant Klebsiella pneumoniae Strain Carrying bla KPC-2 Conjugative Drug-Resistant Plasmid. Front Public Health 2021; 9: 809753.

43 Liu C, Dong N, Zeng Y, et al. Co-transfer of last-line antibiotic resistance and virulence operons by an IncFIBk-FII-X3-ColKP3 hybrid plasmid in Klebsiella pneumoniae. J Antimicrob Chemother 2022; 77: 1856–61.

44 Rawle R, Saley TC, Kang Y-S, et al. Introducing the ArsR-Regulated Arsenic Stimulon. Front Microbiol 2021; 12: 630562.

45 Robertson J, Bessonov K, Schonfeld J, Nash JHE. Universal whole-sequence-based plasmid typing and its utility to prediction of host range and epidemiological surveillance. Microb Genom 2020; 6. DOI:10.1099/mgen.0.000435.

46 Tian C, Shi Y, Ren L, et al. Emergence of IS26-mediated pLVPK-like virulence and NDM-1 conjugative fusion plasmid in hypervirulent carbapenem-resistant Klebsiella pneumoniae. Infect Genet Evol 2023; 113: 105471.

47 Zou H, Zhou Z, Berglund B, et al. Persistent transmission of carbapenem-resistant, hypervirulent Klebsiella pneumoniae between a hospital and urban aquatic environments. Water Res 2023; 242: 120263.

48 Li Y, Zhang P, Du P, et al. Insertion sequences mediate clinical ST34 monophasic Salmonella enterica serovar Typhimurium plasmid polymorphism. Microbiol Res 2023; 272: 127387.

49 Li Y, Wang Q, Peng K, et al. Distribution and genomic characterization of tigecycline-resistant tet(X4)-positive Escherichia coli of swine farm origin. Microb Genom 2021; 7. DOI:10.1099/mgen.0.000667.

50 Bolger AM, Lohse M, Usadel B. Trimmomatic: a flexible trimmer for Illumina sequence data. Bioinformatics 2014; 30: 2114–20.

51 Bankevich A, Nurk S, Antipov D, et al. SPAdes: a new genome assembly algorithm and its applications to single-cell sequencing. J Comput Biol 2012; 19: 455–77.

52 Srijan A, Margulieux KR, Ruekit S, et al. Genomic Characterization of Nonclonal mcr-1-Positive Multidrug-Resistant Klebsiella pneumoniae from Clinical Samples in Thailand. Microb Drug Resist 2018; 24: 403–10.

## References

1 Bolger AM, Lohse M, Usadel B. Trimmomatic: a flexible trimmer for Illumina sequence data. Bioinformatics 2014; 30: 2114–20.

2 Bankevich A, Nurk S, Antipov D, et al. SPAdes: a new genome assembly algorithm and its applications to single-cell sequencing. J Comput Biol 2012; 19: 455–77.

3 Katz LS, Griswold T, Morrison SS, et al. Mashtree: a rapid comparison of whole genome sequence files. J Open Source Softw 2019; 4. DOI:10.21105/joss.01762.

4 Price MN, Dehal PS, Arkin AP. FastTree: computing large minimum evolution trees with profiles instead of a distance matrix. Mol Biol Evol 2009; 26: 1641–50.

5 Price MN, Dehal PS, Arkin AP. FastTree 2--approximately maximum-likelihood trees for large alignments. PLoS One 2010; 5: e9490.

6 Page AJ, Cummins CA, Hunt M, et al. Roary: rapid large-scale prokaryote pan genome analysis. Bioinformatics 2015; 31: 3691–3.

7 Robertson J, Nash JHE. MOB-suite: software tools for clustering, reconstruction and typing of plasmids from draft assemblies. Microb Genom 2018; 4. DOI:10.1099/mgen.0.000206.

8 Argimón S, Abudahab K, Goater RJE, et al. Microreact: visualizing and sharing data for genomic epidemiology and phylogeography. Microb Genom 2016; 2: e000093.

9 Arredondo-Alonso S, Gladstone RA, Pöntinen AK, et al. Mge-cluster: a reference-free approach for typing bacterial plasmids. NAR Genom Bioinform 2023; 5: lqad066.

10 Lam MMC, Wyres KL, Judd LM, et al. Tracking key virulence loci encoding aerobactin and salmochelin siderophore synthesis in Klebsiella pneumoniae. Genome Med 2018; 10: 77.

11 Srijan A, Margulieux KR, Ruekit S, et al. Genomic Characterization of Nonclonal mcr-1-Positive Multidrug-Resistant Klebsiella pneumoniae from Clinical Samples in Thailand. Microb Drug Resist 2018; 24: 403–10.

